# Assessing methods for geometric distortion compensation in 7T gradient echo fMRI data

**DOI:** 10.1101/2020.07.02.184515

**Authors:** Michael-Paul Schallmo, Kimberly B. Weldon, Philip C. Burton, Scott R. Sponheim, Cheryl A. Olman

## Abstract

Echo planar imaging (EPI) is widely used in functional and diffusion-weighted MRI, but suffers from significant geometric distortions in the phase encoding direction caused by inhomogeneities in the static magnetic field (B_0_). This is a particular challenge for EPI at very high field (≥ 7T), as distortion increases with higher field strength. A number of techniques for distortion correction exist, including those based on B_0_ field mapping and acquiring EPI scans with opposite phase encoding directions. However, few quantitative comparisons of distortion compensation methods have been performed using human EPI data, especially at very high field. Here, we compared distortion compensation using B_0_ field maps and opposite phase encoding scans in two different software packages (FSL and AFNI) applied to 7T gradient echo (GE) EPI data from 31 human participants. We assessed distortion compensation quality by quantifying alignment to anatomical reference scans using Dice coefficients and mutual information. Performance between FSL and AFNI was equivalent. In our whole-brain analyses, we found superior distortion compensation using GE scans with opposite phase encoding directions, versus B_0_ field maps or spin echo (SE) opposite phase encoding scans. However, SE performed better when analyses were limited to ventromedial prefrontal cortex, a region with substantial dropout. Matching the type of opposite phase encoding scans to the EPI data being corrected (e.g., SE-to-SE) also yielded better distortion correction. While the ideal distortion compensation approach likely varies depending on methodological differences across experiments, this study provides a framework for quantitative comparison of different distortion compensation methods.

## Introduction

Geometric fidelity is critical for high quality brain imaging. It is essential for accurate interpretation of functional MRI (fMRI) data based on anatomical landmarks and is necessary for precise quantification of structural and functional connectivity. It is also relevant for clinical brain imaging applications, such as neurosurgery and the placement of deep brain stimulation electrodes^2–4^. However, currently-popular MRI techniques suffer from a number of common artifacts that degrade spatial fidelity, including gradient nonlinearities and geometric distortion due to static magnetic field (B_0_) inhomogeneity^5, 6^. A number of methods to correct for these geometric artifacts have been established^5–13^. To select an appropriate method for distortion correction, quantitative comparisons between methods are essential.

Echo planar imaging (EPI) is among the most commonly used MRI techniques in human neuroscience. Rapid acquisition times enable studies of functional brain activation (i.e., fMRI; often ≤ 1 s per whole-brain image) and efficient measurement of white matter tractography via diffusion-weighted MRI (dMRI; on the order of 5 s per image). This temporal efficiency comes at the cost of relatively low pixel bandwidth in the phase encoding (PE) direction, which results in severe geometric distortions in regions of B_0_ inhomogeneity^6–8^. Lower bandwidth (i.e., longer effective echo spacing) makes distortion more severe; distortion of some regions in EPI data in the PE direction often reaches 5-10 mm^6^. B_0_ inhomogeneities and the resulting distortions are greatest at the interface of different tissue types (e.g., brain, bone, air) in regions such as the orbitofrontal cortex and temporal lobes. Inhomogeneities also scale linearly with B_0_ field strength, such that geometric distortions can be more severe at 7 Tesla than at 3 Tesla^14^.

A number of methods for minimizing and correcting geometric distortion in EPI data exist. Prospectively, geometric distortion can be limited by reducing B_0_ inhomogeneity via B_0_ shimming, and confirming shim quality during a scan by measuring the linewidth of the water signal. Geometric distortion can also be limited by shortening read-out time. Methods for this include: 1) using multi-shot or segmented EPI^15–17^ (rather than single-shot sequences, at the cost of longer TRs and increased physiological noise sensitivity), 2) using a higher parallel imaging acceleration factor (R; at the cost of reduced signal-to-noise ratio [SNR]), 3) increasing receiver bandwidth (i.e., reducing echo spacing, using stronger or faster head gradients for example, at the cost of reduced SNR), 4) decreasing the field of view in the PE direction (at the cost of reduced spatial coverage)^18^. Although one might be tempted to think that distortion would also be attenuated by reducing the sampling of *k*-space data in the PE direction using partial Fourier approaches, this is not the case because partial Fourier does not change the pixel bandwidth. Finally, it is worth noting that when using spiral acquisition sequences^19^ in place of EPI, B_0_ inhomogeneity produces blurring rather than geometric distortion, which may be preferable for some applications that do not require high spatial specificity.

It is also possible to correct geometric distortion in an EPI dataset retrospectively, which has been shown to improve registration between EPI and T_1_-weighted anatomical data^7^. A number of different methods for retrospective distortion compensation have been introduced, including:

1. B_0_ field mapping by measuring phase differences from two gradient echo (GE) images with different echo times (TEs)^7, 20, 21^,
2. calculating a distortion field based on two EPI scans with opposite PE directions (i.e. forward & reverse, often anterior-posterior and posterior-anterior; hereafter referred to as opposite phase encoding [oppPE] field mapping), for which the geometric distortion will be equal but in opposite directions^22–24^,
3. non-rigid registration (e.g., affine or spline fitting) of the distorted EPI to a minimally distorted anatomical reference^25–29^,
4. mapping the EPI point-spread function^30–33^,
5. methods based on forward and inverse modeling of the distortion^34, 35^,
6. multi-reference scan methods^36^,
7. hybrid methods (e.g., B_0_ or oppPE field mapping plus non-rigid registration)^8, 11, 21^, and
8. dynamic methods for correcting time-varying geometric distortions due to factors such as head movement^14, 35, 37^. Of these, the first two (B_0_ and oppPE field maps) are arguably the most popular and are currently implemented in various forms across many widely used MRI analysis software packages (e.g., FSL^38^, AFNI^39^, SPM^40^, BrainVoyager^41^). Thus, we chose to focus on quantitative comparisons between B_0_ and oppPE field map approaches in the current study.

With regard to oppPE field maps, it has been suggested that spin echo (SE) EPI scans may offer an advantage over GE sequences in mapping the distortion field^12^, as the former minimizes signal dropout from through-slice dephasing due to the 180° refocusing pulse at TE/2. This suggests that a pair of SE EPI scans with opposite PE directions should give a more complete map of field inhomogeneities than a GE oppPE pair. However, this theoretical motivation has not, to our knowledge, been tested empirically. Alternatively, one may consider whether other factors could limit the utility of SE oppPE field maps for the correction of geometric distortion in GE EPI data. For example, there may be increased opportunity for subject motion due to added scan time, as two additional SE scans must be acquired (forward and reverse PE) versus only one additional GE scan (reverse) to correct GE EPI data (which itself can serve as the forward image in the GE oppPE pair). Further, when using SE oppPE field maps to correct distortion in GE data, image contrast differences between SE and GE could lead to incorrect mapping of voxel shifts in regions of local signal compression^10, 23^. Thus, in the current study, we sought to directly and quantitatively compare the performance of different distortion compensation methods that are currently standard in the field, to explore which method would perform best in our GE fMRI dataset, and whether we might find empirical evidence for the theorized superiority of SE oppPE methods^12^.

Previous studies that have compared different methods for geometric distortion compensation have generally focused on data collected at field strengths of 1.5 to 3T, for which geometric distortion may be less extreme as compared to very high field (≥ 7T). The proliferation of very high field imaging methods^42^, due in part to efforts such as the Human Connectome Project^43–47^, makes it increasingly important to achieve effective geometric distortion compensation of high field EPI data. Therefore, in the current study we examined this issue using 7T EPI data that we have collected as part of the Psychosis Human Connectome Project at the University of Minnesota’s Center for Magnetic Resonance Research.

Prior investigations of geometric distortion compensation methods have not, generally, included correction for additional gradient nonlinearities^48^. Gradient nonlinearities are unrelated to distortion due to B_0_ inhomogeneity, are present in all three dimensions (PE, readout, and through-slice), and are sequence independent^5^. These gradient nonlinearities can be on the order of 1-2%^5, 6^ and will vary between scanners due to differences in gradient hardware. Distortions due to gradient nonlinearities may therefore confound efforts to achieve high spatial fidelity in EPI data, and are particularly important to consider when trying to unify datasets acquired on different scanners (e.g., a T_1_ anatomy acquired at 3T, and GE EPI fMRI data acquired at 7T, as in the current study) or during different scanning sessions.

In this study, we compared the methods noted above (i.e., GE and SE oppPE as well as B_0_ field maps) for the correction of geometric distortion due to B_0_ inhomogeneity in GE EPI data collected at very high field (7T), following a separate initial step to correct for distortion due to known gradient nonlinearities. We sought to answer the following question: which current standard distortion compensation method(s) would perform best for our 7T GE fMRI data? This study presents a framework within which to answer this question for a given dataset. We do not intend to prescribe one method as definitively superior over another in all cases, as relative performance is expected to depend on acquisition parameters, order and timing of the acquisition of EPI and field map scans, scanner and radiofrequency coil hardware, the brain regions being examined, and the details of the processing pipeline that is used. Our results suggest that all of the examined methods improved correspondence between GE EPI and T_1_ anatomical data, with the best performance across the whole brain being observed for GE oppPE field map methods in our dataset.

## Methods

### Participants

We recruited 31 participants for the current study from a larger sample as part of the Psychosis Human Connectome Project. This included 12 patients with a diagnosed psychotic disorder (e.g., schizophrenia), 9 first-degree biological relatives of patients with psychosis (i.e., parents, siblings, or children), and 10 healthy controls. Group differences were not examined in this particular study, as a subject’s mental health status was not deemed relevant to the assessment of geometric distortion compensation methods. We chose to study a diverse population (i.e., including patients and controls, not expert MRI subjects) in order to make our results more broadly applicable to the type of MRI data that would be obtained in clinical populations such as adults with psychosis. Subject demographics were as follows: 20 female and 11 male participants, mean age was 45 years (*SD* = 11 years).

Inclusion criteria for the Psychosis Human Connectome Project were as follows: age 18-65 years, English as primary language, the ability to provide informed consent, no legal guardian, no alcohol or drug abuse within the last 2 weeks, no alcohol or drug dependence within the last 6 months, no diagnosed learning disability or IQ less than 70, no current or past central nervous system disease, no history of head injury with skull fracture or loss of consciousness longer than 30 min, no electroconvulsive therapy within the last year, no tardive dyskinesia, no visual or hearing impairment, no condition that would inhibit task performance such as paralysis or severe arthritis. All patients had a history of bipolar I, schizophrenia, or schizoaffective disorder and were not adopted. Relatives had a biological parent, sibling, or child with a history of one of these disorders and were not adopted. Controls had no personal or family history (parents, siblings, children) of these disorders. Additional inclusion criteria for this particular study included the ability to fit comfortably within the scanner bore (60 cm diameter) and the radio frequency head coil (head circumference less than 62 cm), weight less than 440 pounds, and corrected Snellen visual acuity of 20/40 or better. Further, all participants had completed two 3T fMRI scanning sessions prior to 7T scanning and did not exceed a limit of 0.5 mm of head motion across greater than 20% of TRs from all 3T fMRI runs (approximately 2 hours of scanning). Finally, participants included in this study had all 7T MRI scans acquired in the prescribed order (see below) and did not exceed a limit of 0.5 mm of head motion on greater than 20% of TRs during 7T fMRI scans (1.25 hours of scanning).

All participants provided written informed consent prior to participation and were compensated for their time. All experimental procedures complied with the regulations for research on human subjects described in the Declaration of Helsinki and were approved by the Institutional Review Board at the University of Minnesota. All subjects were found to have sufficient capacity to provide informed consent, as assessed by the University of California Brief Assessment of Capacity to Consent^49^.

### Experimental protocol

7T MRI data were acquired on a Siemens MAGNETOM scanner (software version: VB17). This scanner was equipped with an 8-kW radio frequency power amplifier and body gradients with 70 mT/m maximum amplitude and 200 T/m/s maximum slew rate. We used a Nova Medical (Wilmington, MA) radio frequency head coil with 1 transmit and 32 receive channels for all 7T MRI data acquisition. Subjects were provided with head padding inside the coil and instructed to minimize head movements during scanning. We placed 5 mm thick dielectric pads (3:1 calcium titanate powder in water) under the neck and beside the temples, as this has been shown to improve transmit B_1_ homogeneity in the cerebellum and temporal lobe regions during 7T MRI^44^.

MRI data were acquired using sequences and scan parameters that followed the original Young Adult Human Connectome Project (HCP)^43–47^. Parameters for the different MR scans are listed in Table 1. Additional scan parameters include a multiband acceleration factor of 5 for GE and SE fMRI, and GRAPPA parallel imaging acceleration factor (R) of 2 for all scans except the B_0_ field map (no acceleration). The delta TE for the B_0_ field map scan was 1.02 ms. Single-band reference scans were acquired with each multi-band EPI scan (GE & SE). Data were acquired with an oblique-axial orientation using Siemens Auto Align to standardize the orientation and positioning of the imaging field of view.

**Table 1.** Scan parameters. Anat. = anatomical, TR = repetition time, TE = echo time, FOV = field of view, iso. = isotropic.

| Scan | Field | TR | TE | Echo spacing | Flip angle | Resolution | Partial Fourier | Slices | FOV (mm) |
| --- | --- | --- | --- | --- | --- | --- | --- | --- | --- |
| GE EPI | 7T | 1000 ms | 22.2 ms | 0.64 ms | 45° | 1.6 mm iso. | 7/8 | 85 | 208 x 208 |
| B<sub>0</sub> field map | 7T | 642 ms | 4.08 / 5.1 ms | - | 32° | 1.6 mm iso. | 6/8 | 85 | 208 x 208 |
| SE EPI | 7T | 3000 ms | 60 ms | 0.64 ms | 90° / 180° | 1.6 mm iso. | 7/8 | 85 | 208 x 208 |
| T<sub>1</sub> anat. | 3T | 2500 ms | 1.81 / 3.6 / 5.39 / 7.18 ms | 11.2 ms | 8° | 0.8 mm iso. | Off (phase) & 6/8 (slice) | 208 | 256 x 256 |
| T<sub>2</sub> anat. | 3T | 3200 ms | 564 ms | 3.86 ms | Variable | 0.8 mm iso. | Allowed (phase) & off (slice) | 208 | 256 x 256 |

7T MRI data in this study were acquired in a fixed scan order:

1. auto-align scout and localizer,
2. GE EPI with posterior-anterior (PA) phase encoding direction (3 TRs; Figure 1B),
3. first GE EPI with anterior-posterior (AP) phase encoding direction (324 TRs; Figure 1A),
4. second AP GE scan (297 TRs),
5. B_0_ field mapping scan (Figure 1D & E),
6. AP SE scan (3 TRs; Figure 1G),
7. PA SE scan (3 TRs; Figure 1H),
8. third AP GE scan (468 TRs).

**Figure 1.**
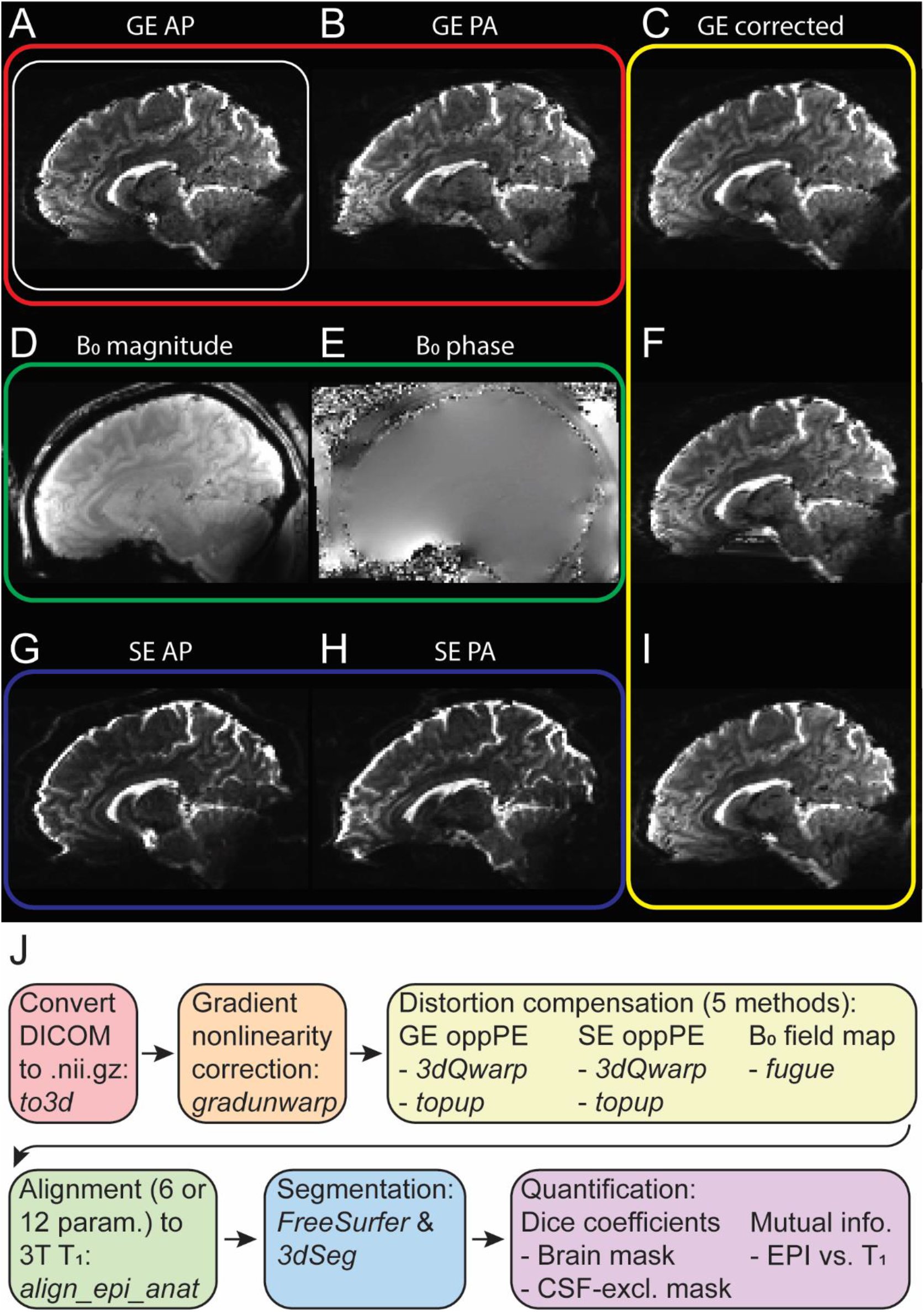
Data and processing pipeline. **A** & **B**) Gradient echo (GE) data with opposite (anterior-posterior [AP] and posterior-anterior [PA], respectively) phase encoding directions. White box in **A** indicates that the GE AP data were the base dataset to which all distortion compensation methods were applied. All brain images are examples from the same parasagittal section in the same subject, after gradient nonlinearity correction has been applied. **C**) GE data corrected for geometric distortion using GE oppPE field map. **D** & **E**) B_0_ field map magnitude and phase data, respectively. **F**) GE data with B_0_ field map distortion compensation applied. **G** & **H**) Spin echo (SE) oppPE data (AP & PA, respectively). **I**) GE data after SE oppPE field map distortion compensation. **J**) A summary of our data processing pipeline steps and software (italics). Arrows indicate the sequence in which processing steps were performed.

Inhomogeneity in the B_0_ field was minimized prior to 7T fMRI data acquisition using the Siemens automated B_0_ shimming procedure (256 × 256 × 256 mm field of view). Shim currents were calculated to minimize field variation within a 130 × 170 × 120 mm^3^ region (i.e., the adjust volume) with an oblique-axial orientation centered on the brain (standardized by Auto Align). To assess shim quality, the linewidth of water (full width at half-maximum [FWHM]) was measured in the Siemens Interactive Shim tab during each scanning session before fMRI data were acquired (for this study, mean linewidth across subjects = 60 Hz, *SD* = 11 Hz). Shim values were stored and applied across all scanning runs using a 3^rd^-party stand-alone program (*shimcache*), to prevent any accidental loss of the B_0_ shim between scans.

3T structural MRI data were acquired on a Siemens MAGNETOM Prisma scanner (software version: VE11C). This scanner was equipped with two RF power amplifiers with a combined power of 40 kW, and body gradients with 80 mT/m maximum amplitude and 200 T/m/s maximum slew rate. Data were acquired using a Siemens 32 channel radio frequency head coil. T_1_- and T_2_-weighted anatomical scans (parameters listed in Table 1) were acquired in the first of two 3T MRI scanning sessions.

### Data analysis and statistics

Our data processing pipeline is summarized in Figure 1J. All data processing steps were performed using either AFNI^39^ (version 18.2.04) or FSL^38^ (version 5.0.9), as noted below. Data were converted from DICOM to compressed (g-zipped) NIFTI format using AFNI’s *to3d* program. We acquired at least 3 TRs for all EPI scans in order to allow the signal to reach steady state. To obtain a single time point for all EPI scans for the sake of computational efficiency, we took the temporal median of 3 TRs at the beginning or end of each scan (i.e., the time points closest to the respective field map scan[s], see below) using AFNI’s *3dTstat*. Averaging was done to improve signal-to-noise; we used the median to prevent non-steady-state signals at the beginning of a scan from biasing the average.

In order to remove known geometric distortions due to gradient nonlinearities, we then performed gradient nonlinearity unwarping using *gradunwarp* (version 1.0.3; github.com/Washington-University/gradunwarp), with the warp field (a.k.a. voxel displacement map) applied using AFNI’s *3dNwarpApply*. This step is comparable to the correction for static gradient nonlinearities available within the HCP pipeline^48^. In Supplemental Figure 1, we show examples of voxel shift maps from gradient nonlinearity unwarping from a single subject. In Supplemental Table 1, we quantify the magnitude of gradient nonlinearity correction across all subjects. This shows that gradient nonlinearity distortion is often small for many voxels within the brain, but can be quite significant (up to about 4 mm) for some voxels. In our typical analysis path (i.e., in other studies), we apply all geometric corrections within a single resampling step to minimize blurring. In the current study, we first applied gradient nonlinearity correction and then separately applied B_0_ inhomogeneity distortion compensation, which allowed us to specifically examine the performance of B_0_ inhomogeneity distortion correction methods implemented in AFNI versus FSL.

#### Distortion compensation

We then performed corrections for geometric distortion due to B_0_ inhomogeneity on our 7T GE EPI data using each of the following five methods:

1. GE opposite phase encoding (oppPE) field map correction^22–24^ via AFNI’s *3dQwarp*.
2. GE oppPE correction with FSL’s *topup*. Distortion correction in each of the two GE oppPE methods was applied to the median of TRs 1-3 from the first AP GE EPI scan (i.e., the scan closest in time to the GE oppPE field maps scans).
3. B_0_ field map correction^7, 20, 21^ in FSL’s *fugue*, applied to the median of TRs 295-297 from the second AP GE scan (the closest to the B_0_ field map scan in time).
4. SE oppPE field map correction using AFNI’s *3dQwarp*.
5. SE oppPE correction with FSL’s *topup*. Distortion correction for both SE oppPE methods was applied to the median of TRs 1-3 from the third AP GE scan (the closest in time to the SE oppPE field maps scans).

Example brain images for GE oppPE correction using *3dQwarp* (Figure 2A) and *topup* (Figure 2B), B_0_ field map correction using *fugue* (Figure 2C), and SE oppPE correction using *3dQwarp* (Figure 2D) and *topup* (Figure 2E) are provided, as well as uncorrected data (Figure 2F). We also present example images showing the difference between GE EPI data before and after distortion compensation in Supplemental Figure 2. The goal of our analysis was to facilitate comparisons of data quality between these five approaches for distortion compensation, which are current standards in the field. The details of each distortion compensation method are provided below (see Code and data availability for a link to our published code for full details). Note that these methods are all designed to correct geometric distortions in the PE direction only (on the order of several millimeters); distortions in the readout and through-slice directions (generally less than 0.1 mm)^20^ are not corrected by these methods and are not considered further in the present study.

**Figure 2.**
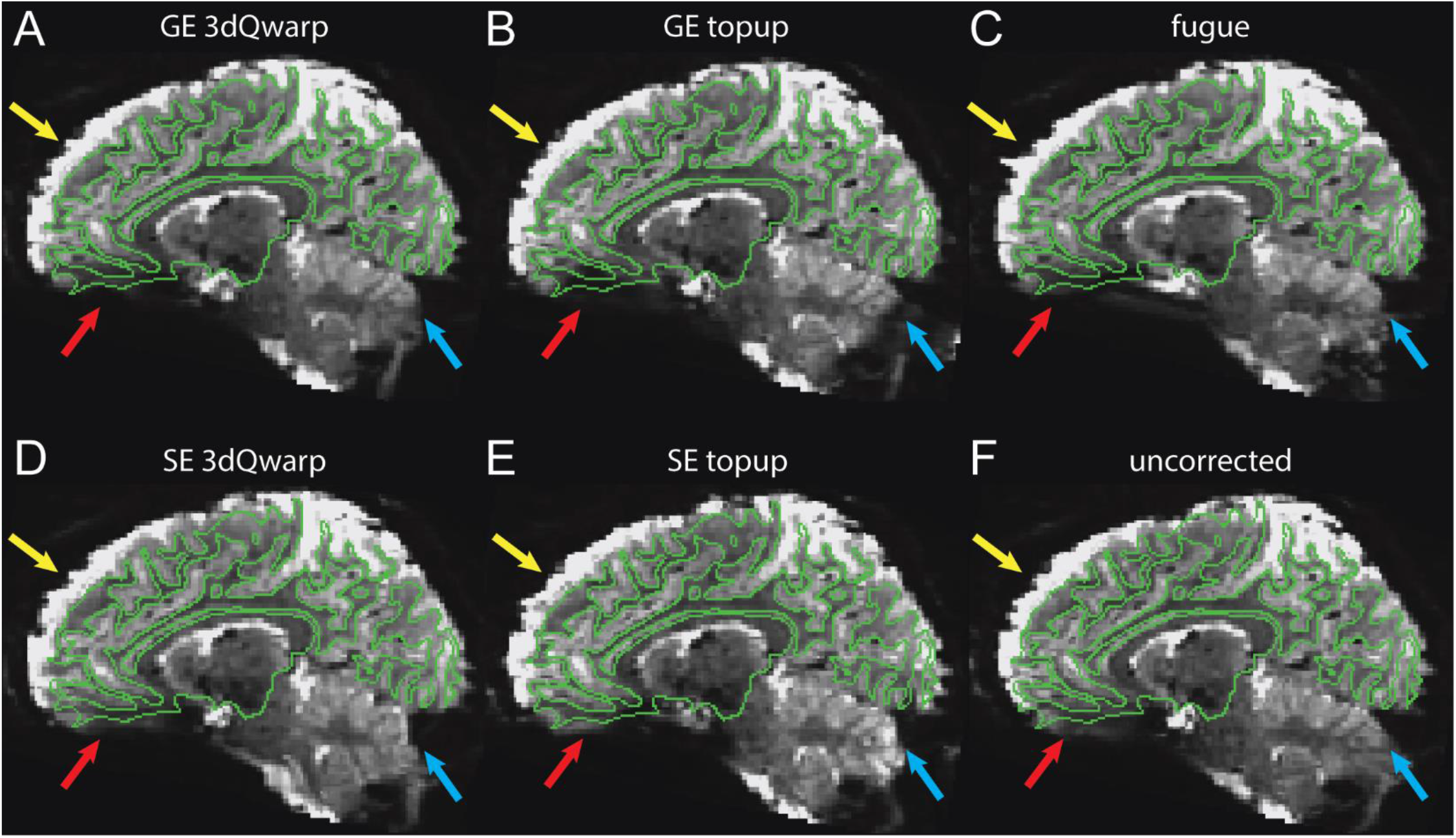
Distortion corrected EPI data. Brain images are shown following distortion compensation using GE oppPE data in AFNI’s 3dQwarp **(A)** or FSL’s *topup* **(B)**, B_0_ field map data via FSL’s *fugue* **(C)**, and SE oppPE data in AFNI’s *3dQwarp* **(D)** or FSL’s *topup* **(E)**, or without distortion correction **(F)**. Green lines show the smoothed white matter surface model from FreeSurfer derived from the T_1_ anatomical data, which illustrates good co-registration with the EPI data overall. Colored arrows indicate regions of interest with notable distortion (**red**: vmPFC, **yellow**: dmPFC, **blue**: posterior regions), which are examined in subsequent analyses. All images show the same sagittal section from the same example subject, after gradient nonlinearity correction and alignment to the T_1_ anatomical data. Note that the EPI data in this image exhibit greater blurring than those in Figure 1C, F, & I (which reflect our true data quality), as the data here have undergone an additional resampling step to transform them into the space of the T^1^ anatomy, to permit visualization of the smoothed white matter surface. Across panels, differences in distortion compensation and alignment are visible between methods.

For the GE and SE oppPE methods using AFNI’s *3dQwarp*, both the AP and PA scans were independently masked using AFNI’s *3dAutomask* to remove non-brain image regions. Next, the warp field for distortion compensation was calculated using *3dQwarp* with the *-plusminus* flag (indicating that the desired undistorted brain image is ‘in between’ the AP and PA scans). This program iteratively calculates the difference in distortion between AP and PA data by fitting cubic polynomials within corresponding three dimensional rectangular image regions of progressively smaller size, starting with the entire volume, down to a minimum region size in our study of 9 mm. Progressive blurring was applied to the patches in each iteration using a spatial median filter. *3dQwarp* allows for progressive blurring to avoid matching fine spatial details in early iterations when attempting to calculate distortion across large image regions. On the first iteration (whole volume), data were median filtered within a radius of 1.6 mm. In subsequent iterations, the median filter radius was set to be 5% of the patch size, such that less blur was applied to smaller image regions. Distortion compensation of the GE EPI scan was then performed by applying the resulting warp field using AFNI’s *3dNwarpApply* with sinc interpolation for the final resampling step. For full details on *3dQwarp*, see afni.nimh.nih.gov/pub/dist/doc/program_help/3dQwarp.html.

For distortion based on GE and SE oppPE field maps using FSL’s *topup*^22^, the warp field was calculated using the default *topup* parameters (i.e., those provided by FSL within the *b02b0.cnf* file). The *topup* method involves calculating the difference in geometric distortion between AP and PA scans across multiple iterations, with progressively finer resolution fits using B-splines. The resolution (knot spacing) for the splines varied across iterations from 19.2 to 3.2 mm in our study. Data were progressively blurred across iterations using a Gaussian kernel with a FWHM ranging from 8 to 0 mm (i.e., no smoothing) for lower to higher resolution fits. Similar to *3dQwarp*, progressive blurring helps to prevent *topup* from matching fine details in early iterations when calculating distortion across larger image regions. Here, data in the first five iterations (i.e., fit with lower resolution) were sub-sampled by a factor of two, to speed up computation time. Prior to *topup*, both the AP and PA scans were zero padded with one additional slice in the superior direction, to obtain an even number of slices (as required for the voxel sub-sampling). Geometric distortion within the GE EPI scan was corrected based on the calculated warp field using FSL’s *applytopup*, with the Jacobian modulation method for intensity restoration and cubic B-spline interpolation. The added empty slice was removed after distortion compensation. This step is analogous to the oppPE method for geometric distortion compensation of fMRI data implemented in the HCP minimal pre-processing pipeline^48^. For more details on *topup*, see fsl.fmrib.ox.ac.uk/fsl/fslwiki/topup/TopupUsersGuide.

For the B_0_ field map method using FSL’s *fugue*, non-brain regions of the magnitude portion of the B_0_ field map were removed using FSL’s *bet*. The difference between the phase portions of the B_0_ field map scans with different echo times were exported by the scanner automatically. This phase difference map was then masked within the extracted brain region, converted from scanner units to radians per second using FSL’s *fsl_prepare_fieldmap* tool (which also includes phase unwrapping), and then blurred (i.e., median filtered, radius = 1.6 mm) using AFNI’s *3dMedianFilter*. This blurring step was done in order to reduce noise in the field map, especially in regions near the outer edge of the brain. Distortion compensation of the GE EPI data was then performed using this phase map via FSL’s *fugue*. For comparison with oppPE methods, we converted phase difference maps to voxel shift maps by multiplying by total readout time and dividing by 2π. We note that unlike *3dQwarp* and *topup*, the *fugue* method produces a voxel shift map in which the values are zero outside of the brain (Figure 5). For full details on *fugue*, please see fsl.fmrib.ox.ac.uk/fsl/fslwiki/FUGUE/Guide.

Our primary analysis (reported in the Analysis #1: main study section of the Results) assumes there was no head motion between each field map and the corresponding GE EPI scan, and that any difference between oppPE scan pairs is caused by geometric distortion and not head motion^7^. Thus, no alignment between field map and GE EPI scans was performed prior to distortion compensation, in order to avoid any spurious ‘correction’ of differences between scans that was in fact caused by geometric distortion (rather than head motion). We performed an additional analysis that included an initial alignment between EPI and field map data, described below as Analysis #2, in order to examine the impact of this methodological decision.

#### Alignment

Following distortion compensation, we aligned GE EPI data and T_1_ anatomical scans using AFNI’s *align_epi_anat.py* function (Figure 2; Figure 3C). This alignment method uses a local Pearson correlation (lpc) cost function^50^, which seeks to co-register brain structures based on correlations between voxels within a small neighborhood (several millimeters). This alignment procedure is similar to boundary-based registration^51^, such as is implemented in the HCP minimal pre-processing pipeline^48^. We performed rigid body alignment (6-parameter: x, y, z, roll, pitch, yaw) to the T_1_ for EPI data from each of the five distortion compensation methods above. We used a rigid body alignment procedure in order to preserve the geometric properties of the GE EPI datasets, to facilitate clear comparison between the various distortion compensation methods. While others, including the original young adult Human Connectome Project^44^, have used 9- or 12-parameter alignment methods to co-register 7T fMRI and 3T anatomical data, in the current study we chose to use a 6-parameter alignment method along with a separate gradient nonlinearity correction step. In this way, we omitted arbitrary scaling of our 7T EPI data to match the 3T anatomy, and instead focused on quantifying the correction of geometric distortions based on first principles.

**Figure 3.**
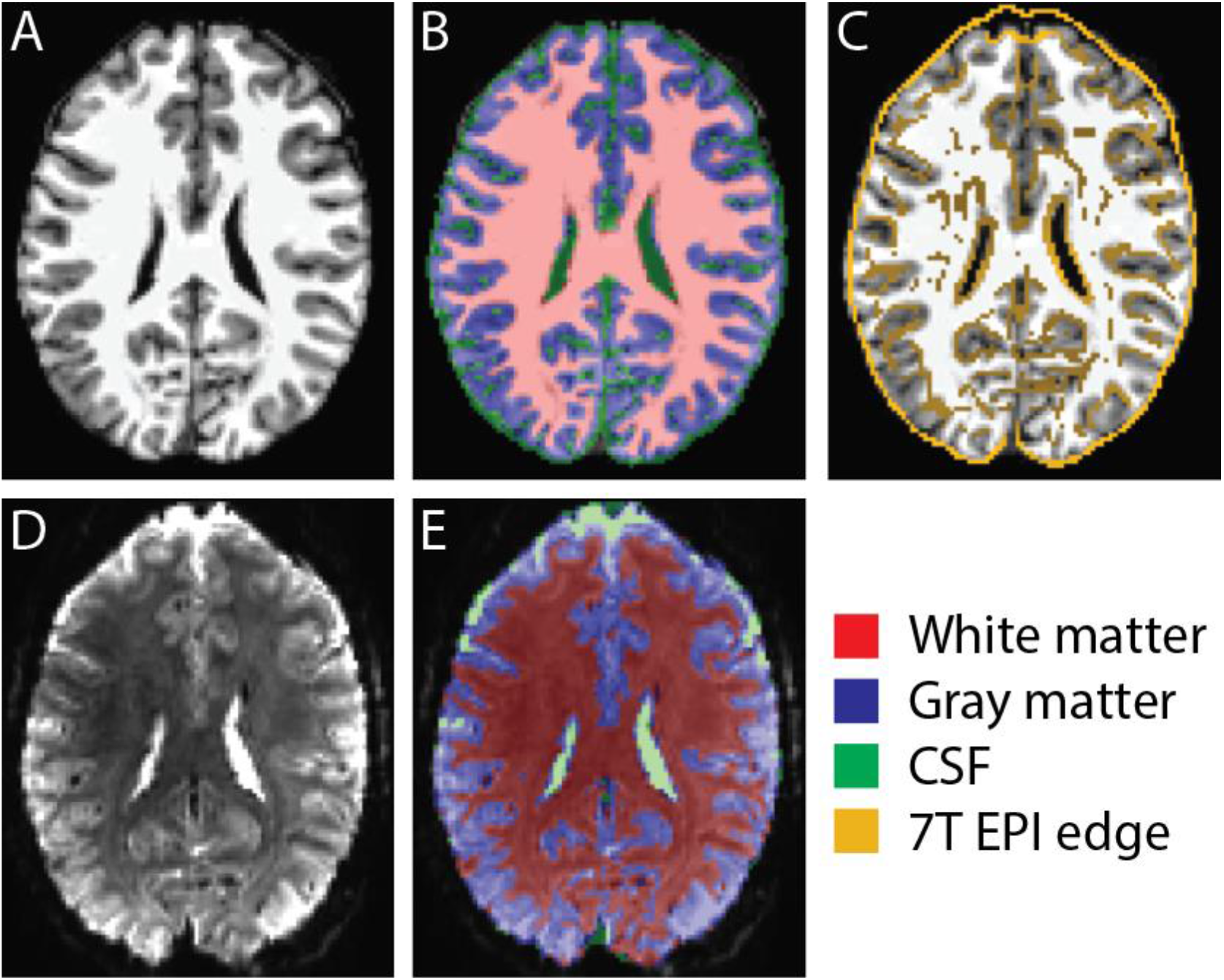
Alignment and segmentation of T_1_ (**A**-**C**) and EPI data (**C**-**E**). Transparent colored overlays in **B** & **E** show binary masks for gray matter (blue), white matter (red), and cerebral spinal fluid (CSF; green) as a result of tissue segmentation. Yellow lines in **C** show edges from 7T GE EPI (using AFNI’s *3dedge3*) overlaid on T_1_ data, to show alignment. All brain images are examples from the same axial section in the same subject, after gradient nonlinearity correction, geometric distortion compensation, and co-registration.

In addition to the five distortion-compensation conditions, we included two further alignment-only analysis conditions in which the GE EPI data were aligned with the anatomical scans without any initial distortion compensation. In the 6-parameter alignment-only condition, EPI data without distortion compensation were aligned with the T_1_ anatomy using 6 parameter (rigid-body) registration with the lpc cost function, as above. In contrast, the 12-parameter alignment only condition used 12 parameter affine alignment (six additional parameters for scaling and shearing). We were motivated to include the 12-parameter alignment method to determine the extent to which the addition of the six scaling and shearing parameters would mirror geometric distortion compensation performed with oppPE or B_0_ field mapping methods.

This procedure yielded a total of seven GE EPI datasets per subject for our analyses (five distortion compensated versions detailed above, plus 6- and 12-parameter alignment-only versions). We refer to these as the seven different analysis conditions below, as they form the basis of our comparison of different approaches for geometric distortion compensation.

Prior to alignment, the T_1_ and T_2_ anatomical data were processed using the HCP minimal pre-processing pipeline^48^ (version 3.22.0), including gradient nonlinearity correction with *gradunwarp* and skull stripping. Note that no correction for geometric distortion due to B_0_ inhomogeneity was performed for these anatomical data, as any such distortions are expected to be minimal (< 0.1 mm)^29^. Although T_2_-weighted scans have a more similar intensity profile to the GE EPI data, T_1_ anatomical scans are currently more widely used in the field of human functional neuroimaging, and robust approaches for aligning EPI and T_1_ data have been developed^50^. Thus, we chose to use the T_1_-weighted scan as our anatomical reference in our primary analysis in order to increase the generalizability of our results. We also conducted an additional analysis (Analysis #4 below) using the T_2_-weighted scan as our anatomical reference, for comparison purposes. We corrected intensity inhomogeneities across the brain in the T_1_ anatomical data using AFNI’s *3dUnifize*.

### Tissue segmentation

In order to facilitate a more detailed analysis of how different distortion compensation methods performed in gray matter and white matter regions, we next segmented different tissue types within our T_1_ and EPI data, with the goal of obtaining brain tissue masks with CSF regions excluded. We performed tissue segmentation for each subject using the T_1_ and T_2_ anatomical scans to define individual white matter and pial surfaces in FreeSurfer^52, 53^ (version 5.3.0) as part of the HCP minimal pre-processing pipeline^48^ (Figure 3A & B). For each of the seven analysis conditions in each subject, we then transformed both the T_1_ and FreeSurfer’s segmentation data (*wmparc*) into the space of the GE EPI scan using the alignment information (obtained above) via AFNI’s *3dAllineate*. Individual binary masks for gray matter and white matter were defined from the transformed *wmparc* file based on FreeSurfer’s anatomical labels (surfer.nmr.mgh.harvard.edu/fswiki/FsTutorial/AnatomicalROI/FreeSurferColorLUT). To define binary masks for cerebral spinal fluid (CSF), we summed the gray and white matter masks, blurred the summed data using AFNI’s *3dmerge* (FWHM = 0.5 mm), and then masked the blurred data at a value of 0.2 to create a binary mask that included the region surrounding the brain (putative CSF). We then summed this mask with a binary mask of the ventricles from FreeSurfer’s *wmparc* file, and subtracted the gray and white matter masks to obtain a CSF mask. Gray matter, white matter, and CSF masks from the T_1_ anatomy were used to aid segmentation of the 7T GE EPI data (below).

To segment the 7T GE EPI data into gray matter, white matter, and CSF regions (Figure 3D & E), we first corrected spatial inhomogeneities in the GE EPI data using AFNI’s *3dUnifize*, and then derived a whole-brain mask using AFNI’s *3dAutomask*. We then segmented the 7T fMRI data from each analysis condition in each subject into gray matter, white matter, and CSF using AFNI’s *3dSeg* function, with the gray matter, white matter, and CSF masks from the T_1_ scan (above) as seed data (*cset* file). Tissue masks from the 7T fMRI data were median filtered with a radius of 1.6 mm using AFNI’s *3dMedianFilter* to reduce noise.

#### Dice coefficients

To quantify alignment between T_1_ anatomical and GE EPI data (and thus the effectiveness of distortion compensation), we calculated the overlap between each subject’s T_1_ and fMRI data in each of the seven analysis conditions using Dice coefficients via AFNI’s *3ddot* function. The Dice coefficient is a measure of how well two binary datasets overlap in three-dimensional space. This metric varies between zero (no overlap) and one (identity) and is calculated by taking the intersection of the two datasets, multiplying by two, and then dividing by the total number of voxels in both scans. Dice coefficients were calculated using two different types of binary masks of the 3T and 7T data: 1) a whole-brain mask (including some CSF) using AFNI’s *3dAutoMask*, and 2) a CSF-excluded mask based on the segmented T_1_ and fMRI data (i.e., including white matter [red] and gray matter [blue], but not CSF [green] regions, as shown in Figure 3B & E). Note that the EPI masks were generated from the distortion corrected EPI data themselves, and not from the corresponding field map scans. We also note that both residual distortion and signal loss due to through-slice dephasing will result in lower Dice coefficient values, indicating poorer agreement with the corresponding T_1_ scan.

#### Mutual information

We also quantified alignment quality by calculating mutual information between the T_1_ and fMRI data for each of the seven analysis conditions in each subject. This metric, which comes from the information theory literature, reflects the similarity of two datasets by quantifying how much is learned about the second dataset from knowing a value in the first. Mutual information is often used to assess multi-modal brain image registration^54^, and should be maximal for two identical datasets that are perfectly aligned. Specifically, mutual information is defined as the difference between the joint entropy and the sum of the marginal entropies for two datasets. Compared to Dice coefficients, mutual information is more sensitive to differences in alignment in internal brain structures. Prior to calculating mutual information, we excluded non-brain regions of the T_1_ and EPI data using the whole-brain masks described above (intensity values for regions outside the mask were set to zero). We computed mutual information using the *mutInfo* function in MATLAB (mathworks.com/matlabcentral/fileexchange/35625-information-theory-toolbox).

### Region of interest (ROI) analyses

To examine the effect of different distortion compensation methods within specific regions of interest (ROIs), we created spherical binary masks in ventromedial prefrontal cortex (vmPFC; radius = 26 mm; Figure 6D), dorsomedial prefrontal cortex (dmPFC; radius = 21 mm; Figure 7D), and a large posterior region encompassing the occipital pole and posterior cerebellum (referred to as the posterior ROI; radius = 34 mm; Supplemental Figure 9). In each subject, we took the intersection of each of these ROIs with the whole-brain masks from the T_1_ and EPI scans for each of the seven analysis conditions to create whole-ROI masks; in Supplemental Figure 3, we show examples of whole-ROI masks from both vmPFC (A & B) and dmPFC (E & F). We also took the intersection between each of the spherical ROIs and the CSF-excluded masks for the T_1_ and EPI data in each condition to define CSF-excluded ROI masks (Supplemental Figure 3C, D, G & H). We then computed the Dice coefficient between these T_1_ and EPI ROI masks, as described for the whole brain above. Finally, we took the intersection between each spherical ROI mask and the T_1_ anatomical and EPI scan data themselves, and then computed mutual information between the T_1_ and EPI data within each ROI for each analysis condition in each subject, excluding non-brain regions as in our whole-brain mutual information calculation above.

#### Additional analyses

We carried out a second analysis (Analysis #2: pre-aligned data) to explore whether differences in head motion across different field map scans may have affected our results. For example, if subjects moved more between and/or during scans that occurred later in the session, then this could have resulted in poorer distortion compensation using certain methods, given that our field map scans were acquired in a fixed order. Whereas our main analysis (above) did not include an initial alignment step between GE EPI data and corresponding field map scans, such an alignment was included in Analysis #2. Specifically, to minimize differences between scans that may be due to head motion, the following scans were aligned to the magnitude portion of the B_0_ field map using a 6-parameter alignment method via AFNI’s *align_epi_anat.py* function: 1) AP GE EPI, 2) PA GE EPI, 3) AP SE EPI, 4) PA SE EPI. In this case, the gradient nonlinearity unwarping (calculated prior to alignment) and the alignment to the B_0_ field map were both applied in a single resampling step using AFNI’s *3dNwarpApply*. Unlike our main analysis above, this procedure assumes that residual head motion between field map scans and fMRI data can be corrected by co-registration to a common reference scan, and that following such an alignment, differences between pairs of field map scans with opposite phase encoding directions reflect geometric distortion due to B_0_ inhomogeneity. Distortion compensation was performed as described above, except that all distortion compensation methods were applied to a single GE EPI scan with AP PE direction (rather than applying distortion compensation to the GE EPI scan acquired closest in time to the corresponding field map scan, as in our main analysis).

In our third analysis (Analysis #3: single-band reference) we examined the role of image contrast in our results by using the single-band reference scan data that were acquired at the beginning of each multi-band 7T GE EPI scan. Image contrast (i.e., white matter vs. gray matter vs. CSF) was higher in the single-band reference as compared to the multi-band data (Supplemental Figure 4). This analysis was identical to the first (main analysis), except that the single-band reference data were used in place of the multi-band 7T GE EPI data during alignment, EPI segmentation, and the quantification of alignment quality using Dice coefficients and mutual information (but were not used to calculate distortion fields). Using the single-band reference data for alignment purposes allowed us to examine the extent to which alignment quality (and thus, our Dice coefficient and mutual information metrics) depended on image contrast. This analysis of the single-band reference data also allowed us to assess whether our EPI segmentation method was limited by image contrast for the multi-band data in our main analysis.

In our fourth analysis (Analysis #4: T_2_ reference anatomy), we investigated whether using the T_2_ anatomical scan (Table 1), rather than the T_1_ as an anatomical reference for alignment might affect our pattern of results. This analysis was otherwise identical to our main analysis, except that we used the lpa cost function (1 – absolute value of the local Pearson correlation)^50^ in AFNI’s *align_epi_anat.py* function, as is recommended for aligning datasets with similar T_2_ weighting.

Our fifth analysis (Analysis #5: posterior-anterior PE data) sought to determine whether our use of GE EPI data with an AP PE direction, versus the opposite PE direction (PA), had an effect of the pattern of results we observed. Here, our analysis was identical to Analysis #2 (pre-aligned data), except that we used the GE PA data as our fMRI scan of interest (to which the various distortion compensation methods were applied), in place of the GE AP data. In this case, we chose to first align all of our data to the magnitude portion of the B_0_ field map scan (as in Analysis #2), since we acquired only a single GE PA scan (see above). We also computed a measure of residual error^33^ by calculating the difference between AP and PA data (from Analyses #2 & 5, respectively), for data from each analysis condition in each subject. In this subtraction analysis, larger intensity differences across voxels indicate greater residual error, and thus poorer distortion compensation.

In our sixth analysis (Analysis #6: Young Adult HCP data) we quantified our distortion compensation metrics as applied to publicly available data from the Young Adult HCP study, in order to facilitate comparisons between available datasets and our own. For more details on the Young Adult HCP study, see ^43, 45–47^. Here, we used unprocessed 7T EPI (fMRI) and 3T T_1_ anatomical data from 20 subjects in the S1200 data release. Gradient nonlinearity correction was not applied. In the first of four conditions in Analysis #6, we applied distortion compensation for each individual’s GE EPI dataset using SE oppPE data via our own *topup* method described above, including 6-parameter (rigid body) alignment of the T_1_ anatomy using AFNI’s *align_epi_anat.py*. We refer to this as the SE *topup* condition. The second (SE *3dQwarp*) condition used AFNI’s *3dQwarp* as described above to apply distortion compensation using the SE oppPE data. In the 6-parameter and 12-parameter alignment-only conditions, GE EPI data (without distortion compensation) were aligned to the T_1_ anatomy via AFNI’s *align_epi_anat.py*, as described for our main analysis above. GE oppPE and B_0_ field maps scans were not available within the 7T Young Adult HCP dataset, so GE *topup*, GE *3dQwarp*, and *fugue* conditions could not be examined in this case. We created whole-brain binary masks from both the EPI and T_1_ data in each of the four analysis conditions using AFNI’s *3dAutomask*. We calculated Dice coefficients and mutual information to quantify T_1_-EPI agreement, as above.

Our seventh analysis (Analysis #7: correcting SE data) examined distortion correction applied to SE, rather than to GE data (as in our other analyses). We chose to examine correction of SE data in order to determine whether matching the data type for the oppPE field map (i.e., SE or GE) to the data to-be corrected, and including the to-be corrected scan as one half of the oppPE scan pair, would affect distortion compensation quality. This analysis was identical to that in Analysis #2 (pre-aligned data), except that the SE scan data (AP PE direction) were used as the EPI data to which all five of the distortion compensation methods and the two alignment-only methods were applied.

All data were visualized in AFNI using default settings for display purposes (i.e., image histograms are scaled so that black ≤ 2%, white ≥ 98%), unless otherwise noted. Brain images are shown in neurological convention (i.e., left is left).

#### Statistics

Statistical analyses were performed in MATLAB (version 2017b). Analyses of variance (ANOVAs) were performed using the *anovan* function, with subjects treated as a random effect, and the seven analysis conditions (i.e., the five different distortion compensation methods, plus the 6- and 12-parameter alignment-only data) as a within-subjects factor. Normality and homogeneity of variance were assessed by visual inspection of the data (e.g., Supplemental Figure 5). Post-hoc comparisons between analysis methods were performed using paired two-tailed *t*-tests, with False Discovery Rate (FDR) correction for twenty-one multiple comparisons (pair-wise comparisons between the seven analysis conditions).

Because the effects of interest (i.e., differences between distortion compensation methods) were within-rather than between-subjects, we used within-subjects error bars^1^ to visualize the variance in each analysis condition, thereby excluding the between-subjects variance for display purposes. To do so, we used an established method^1^ that involved subtracting the mean value for each subject (across all analysis conditions) from all data points for that individual, and then adding the grand mean (across all subjects and conditions).

### Code and data availability

Our analysis code is available on GitHub (github.com/mpschallmo/DistortionCompensation). Imaging data are available from the Human Connectome Project (intradb.humanconnectome.org; first Psychosis Human Connectome Project data release planned for 3^rd^ quarter, 2021).

## Results

### Analysis #1: main study

#### Whole-brain analysis

To compare different distortion compensation methods, we first examined the overlap between whole-brain masks obtained from 7T GE EPI data that had been corrected for geometric distortion and T_1_-weighted anatomical data (T_1_ hereafter), following co-registration (Figure 2; Figure 3C). Data from seven different conditions (i.e., paths to EPI-T_1_ registration) were examined (Figure 1J), including those obtained using five different distortion correction methods, and two alignment-only datasets (6- and 12-parameter alignment). Example brain images corrected using GE oppPE (*3dQwarp* and *topup*), B_0_ field map (*fugue*), and SE oppPE methods (*3dQwarp* and *topup*), as well as uncorrected data, are shown in Figure 2. Binary masks were generated from both the EPI data and the co-registered T_1_ anatomy in each of the seven analysis separately, in each of our 31 subjects. Overlap between EPI and T_1_ masks (Figure 4D) was quantified using the Dice coefficient; higher Dice coefficients reflect more-effective distortion compensation.

**Figure 4.**
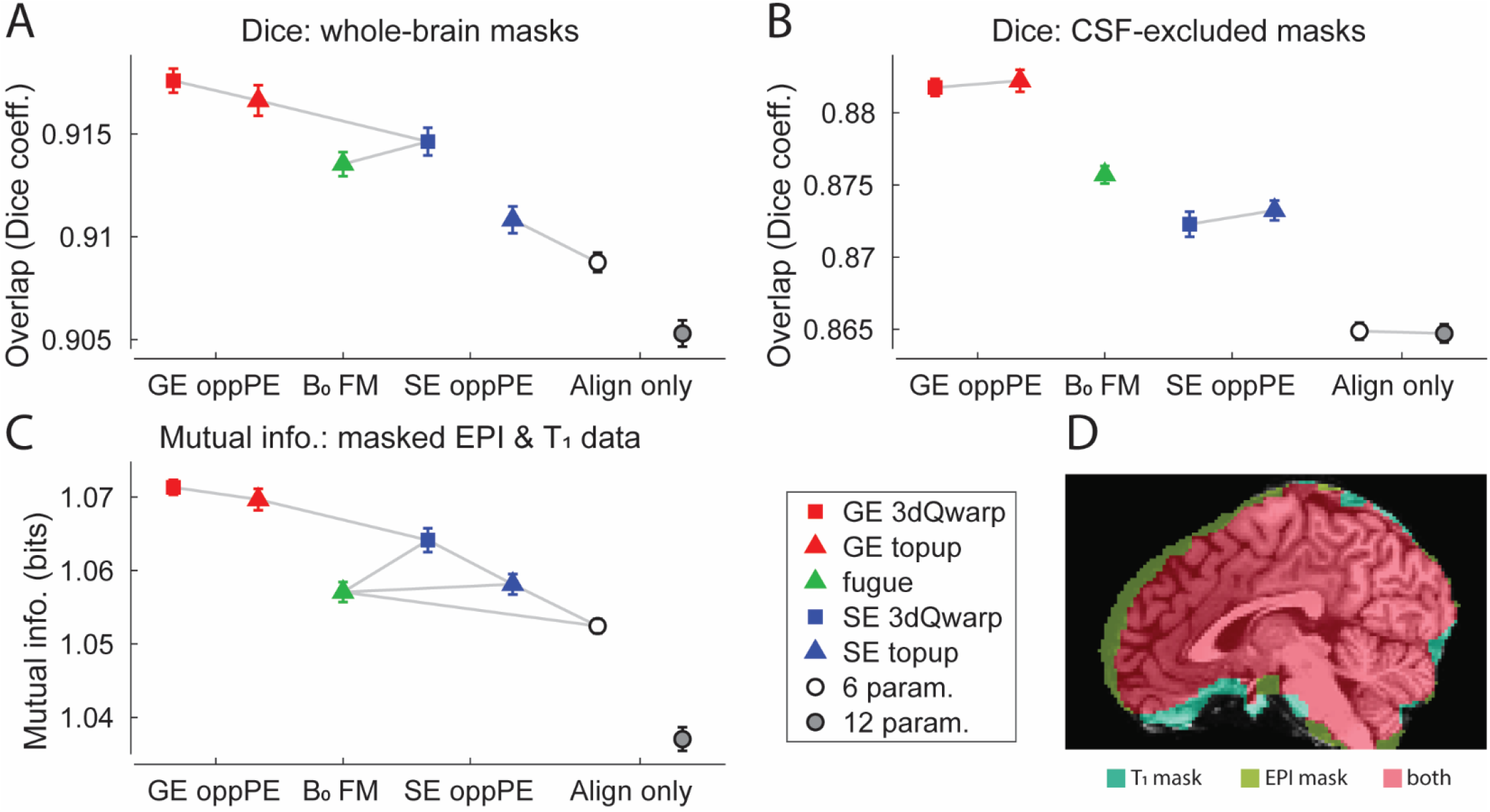
Main whole-brain results. **A)** Overlap (Dice coefficient) between GE EPI and T1 whole-brain masks, across different distortion compensation methods. Gray lines indicate conditions that do <u>not</u> differ significantly (post-hoc paired *t*-tests, threshold p < 0.05, FDR corrected). X-axis labels: GE oppPE = gradient echo opposite phase encoding field map (red), B_0_ FM = B_0_ field map (green), SE oppPE = spin echo opposite phase encoding field map (blue), Align only = alignment-only (no explicit geometric distortion compensation). **B)** Same, but for binary whole-brain masks with regions of cerebrospinal fluid (CSF) excluded, following tissue segmentation. **C)** Mutual information between GE EPI and T_1_ scan data, within the respective whole-brain masks. Squares show data corrected using AFNI, triangles show data from FSL, circles show alignment-only data. Error bars are *SEM* calculated within subjects^1^. **D)** Example T_1_ anatomical image with overlaid whole-brain masks: T_1_ (teal), EPI (yellow), and T1-EPI overlap (red). The Dice coefficient analysis in A quantified the agreement between EPI and T_1_ masks (i.e., red versus teal and yellow), whereas **B** did the same after excluding CSF regions. EPI and T_1_ masks do not overlap perfectly; midline CSF may be included in the automated EPI whole-brain mask (yellow, dorsal), which prompted us to examine CSF-excluded masks **(B)**. Further, through-slice dephasing leads to EPI signal loss in orbital frontal cortex (cyan, ventral), which prompted examination of the vmPFC ROI in Figure 6. All of the tested distortion compensation methods improved agreement between fMRI and T_1_ datasets. For our particular dataset, GE oppPE field maps (red) tended to produce the best results

Dice coefficients for the whole-brain masks were significantly different across analysis conditions (*F*_6,30_ = 42.7, *p* = 4 × 10^−32^), as shown in Figure 4A (see also Supplemental Figure 5 for a visualization of all data points). This indicates that the method of distortion compensation significantly affected the degree to which whole-brain masks from EPI and T_1_ anatomical scans overlapped. Post-hoc paired *t*-tests (FDR corrected for twenty-one comparisons between conditions) revealed that the overlap between EPI and T_1_ anatomical masks was highest and comparable for the two GE oppPE methods using AFNI’s *3dQwarp* and FSL’s *topup* (red symbols). Dice coefficients were lower when using the B_0_ field map (using FSL’s *fugue*; green triangle) and SE oppPE (via *3dQwarp*; blue square). Note that in Figure 4, gray lines indicate conditions that do <u>not</u> differ significantly based on post-hoc tests (i.e., conditions that <u>*do*</u> differ significantly are <u>*not*</u> linked by gray lines; all significant paired *t*_30_ values ≥ 2.82, FDR-corrected *p*-values ≤ 0.042). Lower Dice coefficients were observed for the SE oppPE data corrected using *topup* (blue triangle), which did not differ from the uncorrected data using only a 6-parameter alignment (white circle). Dice coefficients for whole-brain masks were lowest for the data using a 12-parameter alignment only (gray circle).

Next, we examined overlap of tissue masks with CSF regions excluded (see Methods), in order to more closely examine how different distortion compensation methods performed in regions of gray matter and white matter. Dice coefficients for CSF-excluded masks differed significantly between different analysis conditions (*F*_6,30_ = 91.5, *p* = 5 × 10^−52^; Figure 4B), showing that the agreement between non-CSF brain regions from the EPI and T_1_ anatomical scans depended on the method of distortion compensation that was used. Post-hoc paired *t*-tests revealed that the overlap for non-CSF regions was highest when using GE oppPE field maps for distortion compensation (via either AFNI’s *3dQwarp* or FSL’s *topup*; red symbols; all significant paired *t*_30_ values ≥ 2.84, FDR-corrected *p*-values ≤ 0.032). The overlap for the CSF-excluded masks was lower when distortion compensation was performed using a B_0_ field map (via FSL’s *fugue*; green triangle) or SE oppPE field map (in either AFNI’s *3dQwarp* or FSL’s *topup*; blue symbols), and lowest for the alignment-only data (using either 6-parameter [white circle] or 12-parameter alignment methods [gray circle]).

We further compared distortion compensation methods by calculating the mutual information between fMRI and T_1_ data across the whole brain, as higher mutual information reflects better alignment. Compared to the Dice coefficient, mutual information is more sensitive to differences in the alignment of internal brain structures, as it is based on the intensity of all voxels within the brain. Within the whole-brain masks, mutual information between EPI and T_1_ anatomical scans was significantly different across analysis conditions (*F*_6,30_ = 35.1, *p* = 7 × 10^−28^; Figure 4C), reflecting a difference in alignment quality for different distortion compensation methods. In particular, post-hoc tests revealed that mutual information across the brain was generally highest when using the GE oppPE field map methods for distortion compensation (red squares; all significant paired *t*_30_ values ≥ 3.48, FDR-corrected *p*-values ≤ 0.017). Mutual information was generally comparable between B_0_ field map, SE oppPE methods and the 6-parameter alignment-only data (white circle). The 12-parameter alignment-only method (gray circle) yielded lower mutual information compared to all other conditions.

#### Region of interest (ROI) analyses

Thus far, our analyses have focused on metrics to quantify distortion compensation across the whole brain. However, there is significant variability in the amount of geometric distortion due to B_0_ inhomogeneity (and in the voxel shift applied by different correction methods) between brain regions, as shown in Figure 5 and Supplemental Figure 6. Such regional variability is quantified in Supplemental Figure 7, which shows example histograms of voxel shift data obtained using different distortion compensation methods across brain regions from a single subject. Further, Supplemental Figure 8 summarizes the voxel shift data (mean, median, standard deviation, and maximum absolute shift) from different distortion compensation conditions and different brain regions across all subjects.

**Figure 5.**
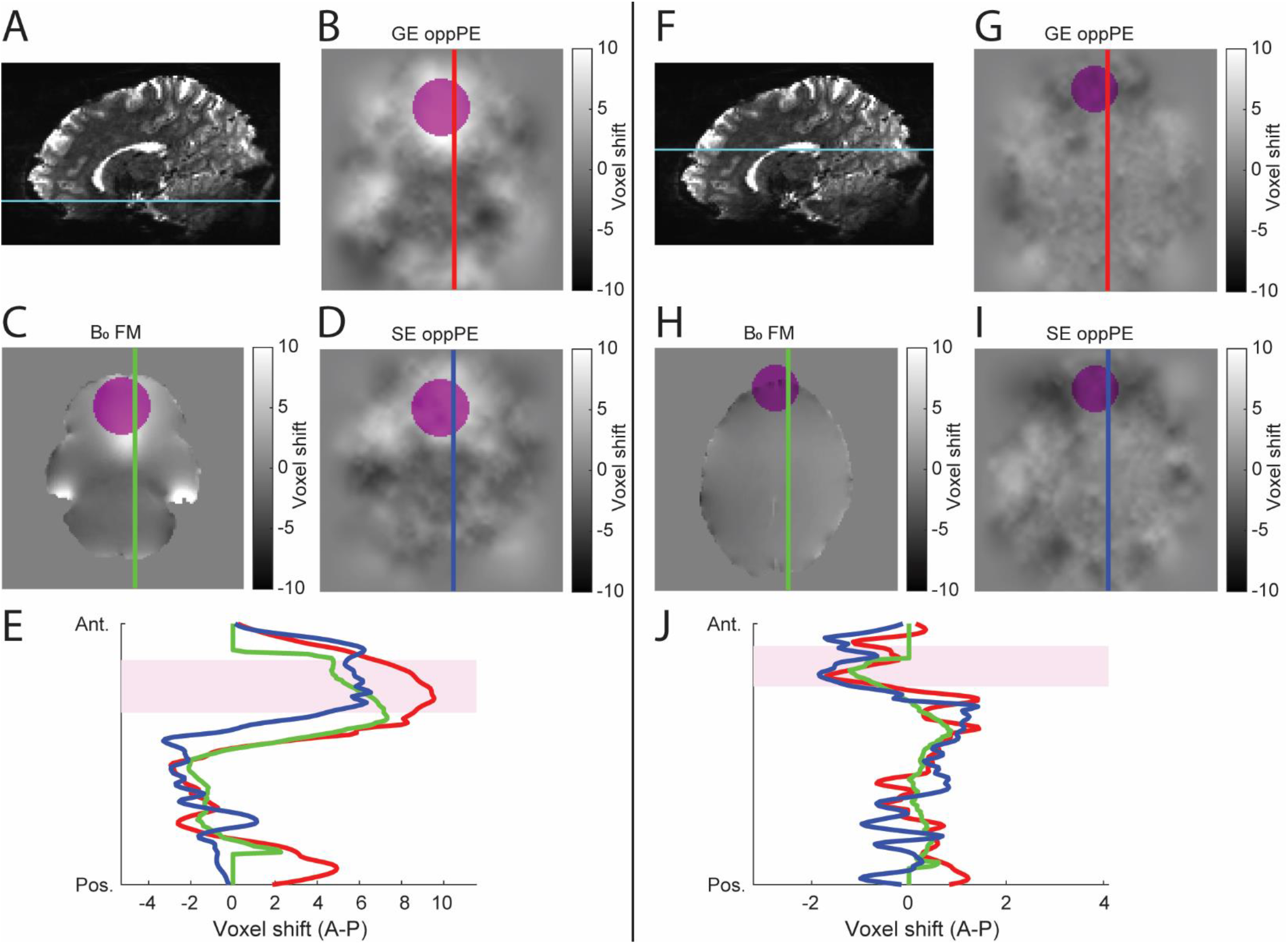
Voxel shift maps for ventral and dorsal regions. **A-E:** ventral regions. Example voxel shift maps from the same ventral axial section (cyan line; **A**) in the same subject are shown for distortion compensation based on GE oppPE (**B**), B_0_ field map (**C**), and SE oppPE (**D**) correction methods. Color bar indicates voxel shift in the anterior-posterior (positive-negative) direction. Colored vertical lines indicate the positions of the voxel shift data plotted in **E**. Magenta regions in **B-E** show the ventromedial prefrontal cortex ROI used below (Figure 6). **F-J**) Same as in **A-E**, but for a more dorsal section. Magenta regions in **G-J** show the dorsomedial prefrontal cortex ROI used below (Figure 7). GE and SE oppPE corrections in these examples were performed with AFNI’s *3dQwarp*. Note that the x-axes differ in **E** and **J**. Differences in voxel shifts between correction methods are apparent, particularly in regions such as ventromedial and dorsomedial prefrontal cortex.

To examine whether our results might be affected by regional variability in distortion due to B_0_ inhomogeneity across the brain, we next compared the performance of different distortion compensation methods within specific regions of interest (ROIs); specifically, the ventromedial prefrontal cortex (vmPFC; Figure 6D, Supplemental Figure 3A & B), the dorsomedial prefrontal cortex (dmPFC; Figure 7D, Supplemental Figure 3C & D), and a posterior ROI which included the occipital pole and posterior cerebellum (Supplemental Figure 6 & Supplemental Figure 9). Each of these ROIs exhibit substantial geometric distortion for EPI scans with an AP PE direction, but in vmPFC there is also significant signal dropout due to through-slice dephasing (Figure 2 & Figure 5A). Thus, we examined the same distortion compensation metrics used above (Dice coefficients and mutual information) but restricted to these ROIs, to assess how different distortion compensation methods may perform in regions with significant distortion, both with and without additional dropout.

**Figure 6.**
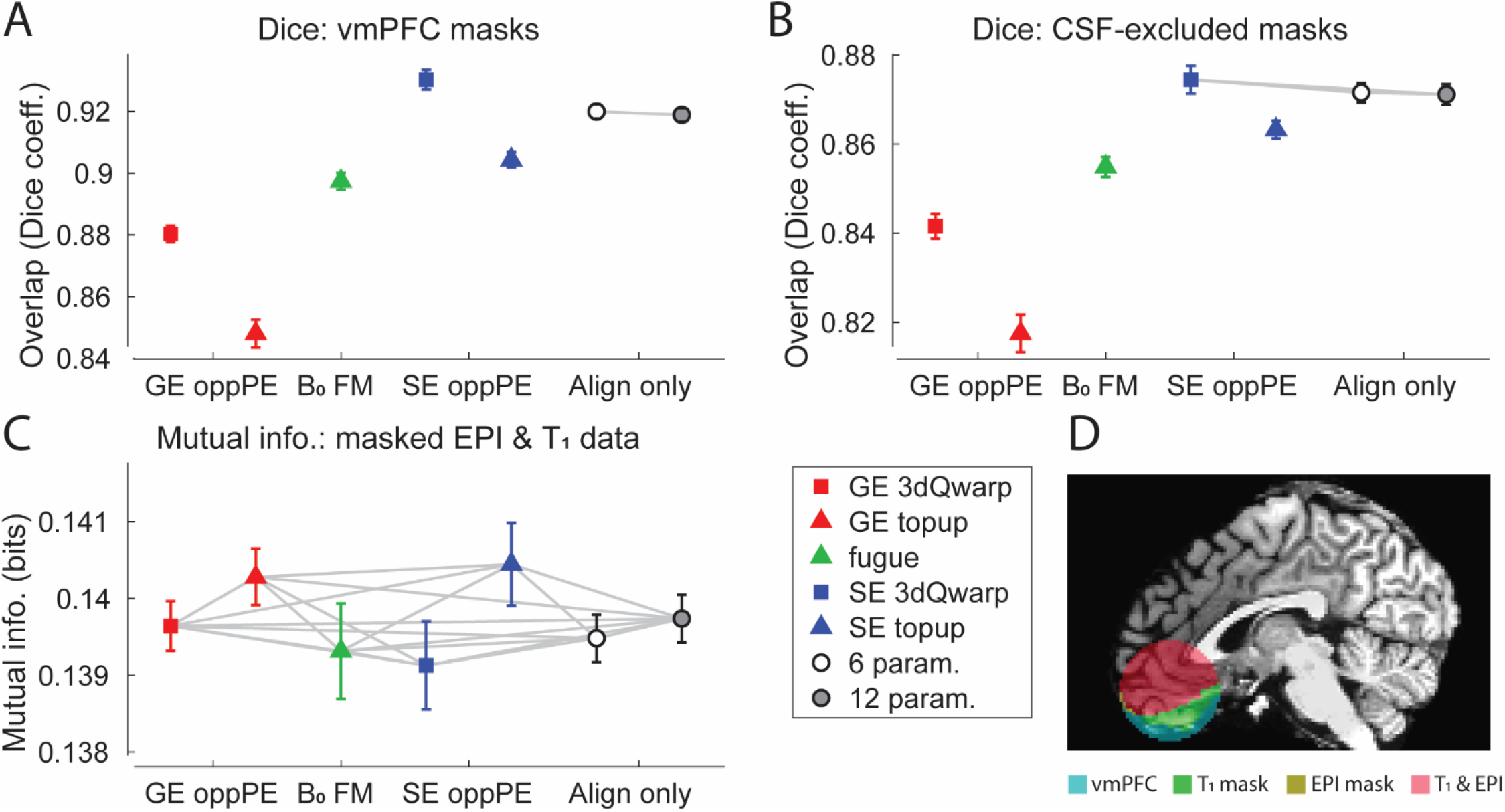
Results in ventromedial prefrontal cortex (vmPFC). **A)** Overlap (Dice coefficient) between GE EPI and T_1_ binary masks restricted to vmPFC, across different distortion compensation methods. Gray lines indicate conditions that do <u>not</u> differ significantly (post-hoc paired t-tests, threshold *p* < 0.05, FDR corrected). X-axis labels: GE oppPE = gradient echo opposite phase encoding field map (red), B_0_ FM = B_0_ field map (green), SE oppPE = spin echo opposite phase encoding field map (blue), Align only = alignment-only (no explicit geometric distortion compensation). **B**) Same, but for binary masks in vmPFC with regions of cerebrospinal fluid (CSF) excluded, following tissue segmentation. **C**) Mutual information between GE EPI and T_1_ scan data in the vmPFC ROI. Squares show data corrected using AFNI, triangles show data from FSL, circles show alignment-only data. Error bars are *SEM* calculated within subjects^1^. **D**) Example T_1_ anatomical image with overlaid vmPFC masks: whole-ROI (teal), T_1_ (green), EPI (yellow), and T_1_-EPI overlap (red). The Dice coefficient analysis in **A** quantified the agreement between EPI and T_1_ masks (i.e., red versus green and yellow), whereas **B** did the same after excluding CSF regions. SE oppPE methods yielded somewhat better T_1_-EPI agreement than GE or B_0_ methods for whole-vmPFC masks (**A**) and CSF-excluded masks in vmPFC (**B**), but little or no advantage over alignment-only methods (gray and white), and there were no significant differences in mutual information across methods (**C**). This suggests a modest advantage for SE oppPE correction over other methods within regions with substantial dropout, but even SE oppPE correction may not yield substantially better T_1_-EPI agreement as compared to uncorrected EPI data (i.e., alignment only).

**Figure 7.**
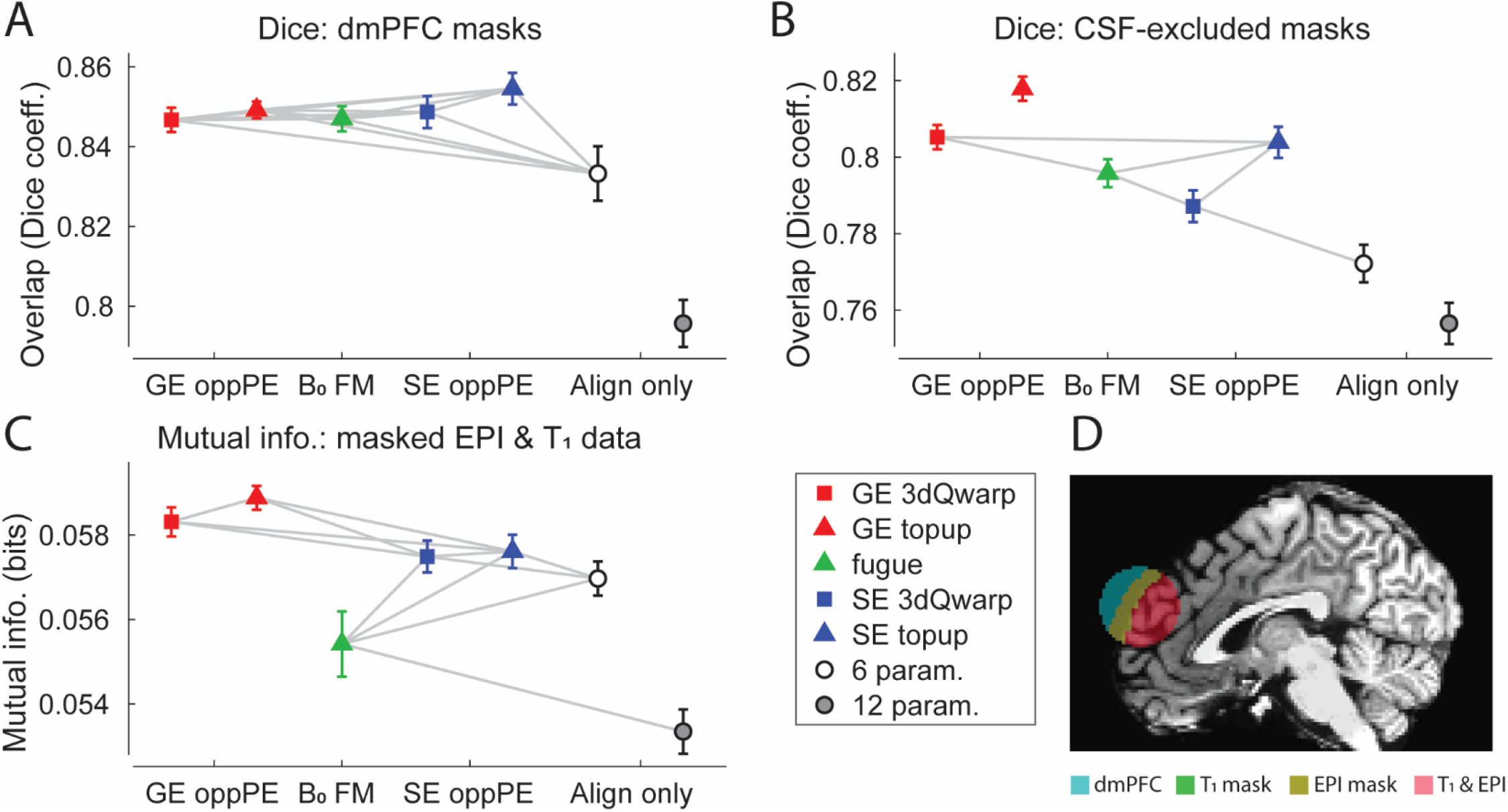
Results in dorsomedial prefrontal cortex (dmPFC). **A**) Overlap (Dice coefficient) between GE EPI and T_1_ binary masks restricted to dmPFC, across different distortion compensation methods. Gray lines indicate conditions that do <u>not</u> differ significantly (post-hoc paired t-tests, threshold *p* < 0.05, FDR corrected). X-axis labels: GE oppPE = gradient echo opposite phase encoding field map (red), B_0_ FM = B_0_ field map (green), SE oppPE = spin echo opposite phase encoding field map (blue), Align only = alignment-only (no explicit geometric distortion compensation). **B**) Same, but for binary masks in dmPFC with regions of cerebrospinal fluid (CSF) excluded, following tissue segmentation. **C**) Mutual information between GE EPI and T_1_ scan data in the dmPFC ROI. Squares show data corrected using AFNI, triangles show data from FSL, circles show alignment-only data. Error bars are *SEM* calculated within subjects^1^. **D**) Example T_1_ anatomical image with overlaid dmPFC masks: whole-ROI (teal), T_1_ (green), EPI (yellow), and T_1_-EPI overlap (red). The Dice coefficient analysis in **A** quantified the agreement between EPI and T_1_ masks (i.e., red versus green and yellow), whereas **B** did the same after excluding CSF regions. T_1_-EPI agreement was generally highest in dmPFC for EPI data corrected using GE oppPE field maps (red) similar to our whole-brain results (Figure 4).

We first asked how different distortion compensation methods would perform in vmPFC (Figure 6D, Supplemental Figure 3A & B), a region that shows substantial geometric distortion and dropout. In particular, we were interested in whether SE oppPE field maps might outperform GE oppPE methods, as superior distortion correction for SE field maps has been theorized on the basis that the 180° refocusing pulse reduces dropout in SE sequences^12^. Dice coefficients for vmPFC masks (Figure 6A & B) showed higher overlap between T_1_ anatomical and EPI data for SE oppPE field map methods (blue) versus GE oppPE (red), with B_0_ field map correction using *fugue* (green) in the middle (ANOVAs, main effects of condition for whole-vmPFC masks and CSF-excluded masks, *F*_6,30_ > 46.7, *p*-values < 3 × 10^−34^). However, there was little or no advantage for any of these methods when compared to scans with no distortion compensation applied at all (6- or 12-parameter alignment only; white and gray, respectively). This suggests that although SE oppPE methods yielded the best agreement between T_1_ and EPI data in the vmPFC region, distortion compensation may not substantially improve T_1_-EPI agreement in vmPFC, compared to no correction. Further, mutual information between T_1_ and EPI data in vmPFC was not significantly different across conditions (ANOVA, main effect, *F*_6,30_ = 0.99, *p* = 0.4), suggesting that none of the distortion compensation methods we tested had a substantial effect on T_1_-EPI agreement in vmPFC according to this metric.

Next, we compared the performance of different distortion compensation methods within a region that also shows substantial geometric distortion due to B_0_ inhomogeneities, but less dropout due to through-slice dephasing, dmPFC (Figure 7D, Supplemental Figure 3C & D). Here, we found results that were more similar to our whole-brain results, as compared to those from vmPFC. In particular, we saw substantial differences in dmPFC across distortion compensation methods for all three of our metrics (ANOVAs, main effects of condition for whole-dmPFC masks, CSF-excluded masks, and mutual information, *F*_6,30_ > 14.1, *p*-values < 4 × 10^−13^; Figure 7), with GE oppPE methods (red) generally yielding the best agreement between T_1_ and EPI data in dmPFC. Thus, we conclude that in regions such as dmPFC in which there is significant geometric distortion (but not dropout), GE oppPE methods may show slight advantages over other distortion compensation techniques, similar to the pattern of results from our earlier whole-brain analysis.

Finally, we examined how distortion compensation varied in posterior brain regions across our seven analysis conditions (Supplemental Figure 6 & Supplemental Figure 9). Like dmPFC, posterior regions exhibit geometric distortion due to B_0_ inhomogeneity, but less through-slice dephasing than in vmPFC. Results within the posterior ROI differed significantly across condition (ANOVAs, main effects of condition for whole-ROI masks, CSF-excluded masks, and mutual information, *F*_6,30_ > 10.1, *p*-values < 1 × 10^−9^), and were generally similar to those in dmPFC. Agreement between T_1_ and EPI data in posterior regions tended to be higher in the five GE oppPE, B_0_ field map, and SE oppPE conditions, as compared to the alignment-only conditions, but results were similar between the five distortion compensation conditions overall.

### Analysis #2: pre-aligned data

Human subjects, especially those who are not experienced with MR scanning, may be more likely to move, or move more toward the end of a long scanning session. Because our scans were acquired in a fixed order, we considered whether differences in head motion might have biased our results in favor of the GE oppPE data, which was acquired near the beginning of the session (approximately 1.25 hours in total length), rather than the B_0_ field map or SE oppPE data, which were acquired near the end. Specifically, if subjects tended to move more during the B_0_ field map and SE oppPE scans, or moved more between these scans and the GE EPI scans on which distortion compensation was performed, then this might degrade the quality of distortion compensation for the B_0_ field map and SE oppPE methods as compared to the GE oppPE method.

To explore this issue, we performed our analyses again after first aligning all field map and 7T GE fMRI scans to the magnitude portion of the B_0_ field map (see Methods for details, and a discussion of why this initial alignment step was omitted from the first analysis). We applied all five distortion compensation methods to the same 7T GE EPI scan, to mitigate any possible bias caused by the fixed scanning order. We found that whether or not the GE EPI and field map data were initially aligned had very little impact on our results across the whole brain (Supplemental Figure 10). After initial alignment, all distortion compensation methods improved agreement between fMRI and T_1_ data (ANOVAs, main effects of condition for whole-brain masks, CSF-excluded masks, and mutual information, *F*_6,30_ > 35.7, *p*-values < 3 × 10^−28^) as compared to the alignment-only conditions, with the GE oppPE field maps showing the strongest performance, similar to the results in our main analysis (Figure 4).

### Analysis #3: single-band reference

Alignment and segmentation of GE EPI data may depend on image contrast (e.g., gray matter vs. white matter intensity). To explore the role of image contrast in our results, we repeated our main whole-brain analyses using the single-band reference data in place of the multi-band 7T GE EPI data for alignment, segmentation, and quantification purposes, as image contrast was higher in the single-band reference (Supplemental Figure 4). The patterns of results for the single-band reference data (ANOVAs, main effects of condition, *F*_6,30_ > 35.2, *p*-values < 5 × 10^−28^; Supplemental Figure 11) were very similar to those obtained with multi-band GE EPI in the main analysis (Figure 4), suggesting that the quality of the alignment and segmentation of our 7T data were not limited by image contrast in the multi-band scans.

### Analysis #4: T_2_ reference anatomy

The alignment between GE EPI data and an anatomical reference, and therefore our subsequent quantification of distortion compensation metrics (Dice coefficients and mutual information), may depend on the chosen anatomical reference scan. To examine this, we repeated our main whole-brain analysis using the T_2_ anatomical data in place of the T_1_. We found that using a T_2_-weighted anatomical scan as a reference for alignment purposes had little effect on our results (ANOVAs, main effects of condition, *F*_6,30_ > 50.4, *p*-values < 5 × 10^−36^; compare results in Supplemental Figure 12 and Figure 4).

### Analysis #5: posterior-anterior PE data

The pattern of geometric distortion due to B_0_ inhomogeneity in EPI data depends critically on the phase encoding (PE) direction (Figure 1)^22–24, 55^. To examine whether our findings were specific to data with an anterior-posterior (AP) PE direction, we repeated an earlier whole-brain analysis (#2: pre-aligned data) using GE EPI data with the opposite (PA) PE direction (see Methods for details). The results of this analysis largely recapitulated our earlier findings (ANOVAs, main effects of condition, *F*_6,30_ > 26.1, *p*-values < 2 × 10^−22^; compare data in Supplemental Figure 13 with those in Supplemental Figure 10), suggesting that the choice of an AP or PA PE direction did not have a dramatic effect on our results. In Supplemental Figure 14 we show the subtraction between distortion-corrected AP and PA data in an example subject. This method has previously been used to quantify residual error, as a metric for distortion compensation quality^33^. As shown in Supplemental Figure 15, across all subjects, residual error for AP – PA data is lowest in the GE oppPE conditions (ANOVA, main effect of condition, *F*_5,30_ = 661, *p* < 9 × 10^−98^), consistent with superior distortion compensation.

### Analysis #6: Young Adult HCP data

To assess how well our findings would generalize outside our own dataset, we conducted another analysis using similar methods with 7T GE EPI data from the Young Adult HCP dataset^43, 45–47^. We examined publicly available datasets from 20 subjects. We assessed data without distortion compensation (6- or 12-parameter alignment only), as well as with SE oppPE distortion correction (using AFNI’s *3dQwarp* or FSL’s *topup*). Example brain images before and after correction are shown in Supplemental Figure 16A & B. Thus, we used the publicly available SE oppPE field maps scans (GE oppPE and B_0_ field map data were not available) to examine SE oppPE correction of the Young Adult HCP data within our own analysis pipeline, in order to assess the effectiveness of this method on an independent dataset.

Agreement between 3T T_1_ anatomical and 7T GE EPI data (Dice coefficients for whole-brain masks and mutual information) from the Young Adult HCP varied significantly across analysis conditions (ANOVA, main effects of condition, *F*_3,19_ > 66.1, *p*-values < 2 × 10^−18^; Supplemental Figure 16C & D). Distortion compensation using SE oppPE field maps tended to improve correspondence between EPI and T_1_ data from the Young Adult HCP (compared to alignment-only data), and yielded similar results to those obtained using SE oppPE methods in our main analysis (compare blue symbols to those in Figure 4).

### Analysis #7: correcting SE data

Up to this point, we have focused on distortion correction of GE EPI data at 7T. Our results have generally shown superior performance for GE oppPE field map scans over SE oppPE methods (e.g., Figure 4), except in regions with severe dropout (i.e., vmPFC; Figure 6). One possibility is that GE oppPE field maps may not be superior overall, but may be better for correcting GE oppPE data in particular, whereas SE oppPE field maps may yield superior correction of SE data. We sought to address this by applying our analyses to SE, rather than GE data. As predicted, distortion compensation quality metrics varied significantly across conditions (ANOVA, main effects of condition, *F*_4,19_ > 153, *p*-values < 2 × 10^−35^; Supplemental Figure 17), with SE oppPE methods outperforming GE oppPE methods when applied to SE EPI data. Alongside our other results, this suggests that matching the oppPE field map to the data type to-be corrected yields superior distortion compensation.

## Discussion

Our analyses showed that all of the distortion compensation methods tested (GE oppPE field maps, B_0_ field maps, SE oppPE field maps) yielded overall improved correspondence between GE fMRI and T_1_ anatomical data across the whole brain, compared to alignment-only data. We found very few differences when comparing our results for oppPE field map corrections performed using AFNI versus FSL (squares vs. triangles, Figure 4), suggesting that these two software packages generally yield equivalent data quality for this type of distortion compensation. However, we did find small but consistent differences in Dice coefficients and mutual information between the various distortion compensation methods we examined. Agreement between GE fMRI and T_1_ data across the whole brain was generally highest in our dataset when using GE oppPE field maps for distortion compensation (red symbols, Figure 4). Hence, we have chosen to implement this particular correction method within our own internal data processing pipeline for the Psychosis Human Connectome Project.

Although it has been theorized that SE oppPE field maps might yield overall better distortion compensation than GE oppPE methods^12^, we generally observed better distortion correction, as quantified by Dice coefficients and mutual information, for GE vs. SE oppPE methods in our whole-brain analyses of GE data (e.g., red vs. blue symbols, Figure 4). However, Analysis #7 (correcting SE data) showed that SE oppPE methods did outperform GE oppPE field maps when applied to correct SE EPI data in particular. Thus, we conclude that matching the type of oppPE field map to the EPI data to-be corrected (whether GE or SE), and including the EPI data of interest as one half of the oppPE pair, yields better distortion compensation than correcting one type of data with the other type of oppPE field map data. We offer two plausible explanations for this finding below.

A similar overall pattern to our whole-brain results (Figure 4) was observed within dmPFC (Figure 7), a region with significant geometric distortion, but not in vmPFC (Figure 6), where both significant distortion and substantial signal dropout due to through-slice dephasing are observed. In vmPFC, we saw somewhat better performance (higher Dice coefficients; Figure 6A & B) for SE versus GE oppPE methods, consistent with the notion that SE oppPE field maps would yield better distortion compensation due to reduced through-slice dephasing^12^. However, Dice coefficients for the SE oppPE methods were generally not higher than for the alignment-only conditions, and we saw no differences in mutual information between T_1_ and EPI data within vmPFC across conditions (Figure 6C). In contrast, the patterns of results in the dmPFC ROI (Figure 7) were similar to those from the whole-brain analyses (Figure 4). Together, these results suggest that for GE EPI data, the advantages (if any) of the SE oppPE approach were limited to regions of the brain with strong drop-out (such as vmPFC), whereas the GE oppPE approach tended to produce better results in our GE data overall.

This study provides a framework for deciding which distortion compensation method to use for a given dataset, based on quantitative comparisons of the agreement between distortion-corrected EPI data and an anatomical reference scan. We expect that the relative performance of different methods may vary across datasets based on data acquisition parameters, scanner and coil hardware, and the details of the processing pipelines that are used. Thus, the reader may wish to compare the relative performance of different distortion compensation methods in their own dataset using an approach similar to ours. We used multiple metrics to quantify EPI-T_1_ agreement as a proxy for correction quality (i.e., Dice coefficients for whole-brain masks and CSF-excluded masks, as well as mutual information), since we acknowledge that there is no single gold standard for measuring the quality of distortion compensation in human brain imaging data^11^ (but see the following studies that used simulations to try to establish ground truth^9, 10^). Across our analyses, we found that certain methodological decisions (i.e., whether or not to align data prior to distortion compensation, the use of single-band reference, T_2_ anatomical, or PA GE data) had little effect on our pattern of results (compare Figure 4 with Supplemental Figure 10, Supplemental Figure 11, Supplemental Figure 12 & Supplemental Figure 13), whereas regional variations in distortion compensation metrics were more dramatic (Figure 6 & Figure 7; Supplemental Figure 9). We found results comparable to our main findings when examining data from the Young Adult HCP study (Supplemental Figure 16). By making our data and analysis code publicly available (see Methods), we hope to facilitate the empirical selection of effective approaches for geometric distortion compensation in future research.

In addition to geometric distortion compensation, our analyses included gradient nonlinearity correction^5, 6, 48^, a post-processing step to correct for static spatial non-uniformities in the brain images caused by the gradients themselves (i.e., not dependent on the scanning sequence or B_0_ field inhomogeneity). In our data, we found gradient nonlinearities led to voxel shifts up to about 4 mm in some regions (Supplemental Figure 1; Supplemental Table 1). Correcting for this type of image distortion is particularly important in cases such as ours, where one wishes to align EPI and anatomical data acquired in different scanning sessions, as distortions due to gradient nonlinearities will vary across scanners based on differences in gradient hardware, and across scanning sessions based on head position. Previous studies comparing different geometric distortion compensation methods have generally not included (or reported) gradient nonlinearity correction. For datasets acquired in a single scanning session, sequence-independent gradient nonlinearities limit geometric fidelity but not the ability to align distortion corrected EPI to anatomical reference scans. We believe that effective corrections for both gradient nonlinearities and geometric distortions are critical for achieving high spatial fidelity, and for harmonizing EPI and T_1_ data across different scanning sessions or field strengths.

We did not initially hypothesize that GE oppPE methods would outperform SE methods in our GE data, but rather sought to explore which method would perform best in our dataset. We offer two possible explanations for this finding, which are not mutually exclusive. First, there may be more opportunities for head motion to degrade the quality of distortion compensation when using a SE oppPE field map to correct GE EPI data (or vice-versa), as there are two additional scans (forward and reverse PE SE) during and between which the subject must hold still, versus only one additional scan for a GE oppPE field map (reverse GE, as the GE EPI data to-be corrected may serve as the forward PE half of the GE oppPE pair). As noted below, any head motion between scans will change the B_0_ inhomogeneities and subsequent geometric distortions, leading to poorer correction. Future studies that acquire multiple SE oppPE field map scans at different time points, or in which subjects are explicitly instructed to move their heads during field maps, may be better able to directly examine this possibility.

Second, superior performance of the GE oppPE field maps for correcting GE data might possibly be attributed to differences in image contrast between GE and SE data^10, 23^ (compare Figure 1A & B vs. Figure 1G & H). In regions of significant B_0_ inhomogeneity, geometric distortion can cause displaced signal from multiple voxels to ‘pile up’ within a single voxel (i.e., local compression)^22, 26, 27, 36^. Local compression depends on PE direction; oppPE field map methods attempt to correct for compression by using interpolation to recreate the image ‘in between’ the forward and reverse PE scans. If local compression differs between SE and GE scans, as would be expected due to contrast differences, then the mapping of spatial information during oppPE distortion compensation may also differ. In this case, the warp field calculated from SE oppPE field maps (and applied to GE EPI data; or vice versa) may be incorrect in regions of local compression, resulting in poorer distortion compensation^10, 23^. This issue of signal compression during distortion compensation has also been appreciated in the dMRI field; local compression differs for scans with different diffusion weighting (and thus different signal contrast), which is relevant for distortion compensation of such data using oppPE methods^10, 23^.

Our results agree with previous studies that have universally shown corrections for geometric distortion due to B_0_ field inhomogeneity improve EPI data quality and alignment with minimally-distorted reference scans^7^. In particular, previous work has generally shown better performance for oppPE field map strategies, as compared to B_0_ field maps, which has been attributed in part to the difficulty of using B_0_ field maps to correct distortion near the edges of the brain (see Figure 1F), where phase values change rapidly across space. Using simulated EPI data, both Esteban^9^ and Graham^10^ showed quantitatively that ground truth undistorted images were recovered best using an oppPE field map method, whereas B_0_ field maps performed slightly worse, and nonlinear registration-based methods were greatly inferior (but still better than no correction at all). Similar conclusions were reached by Hong and colleagues^11^ using SE EPI in the mouse brain at 7T, by Holland and colleagues^12^ using SE EPI at 1.5 and 3T in the human brain, and by Wang and colleagues^13^ using 3T dMRI data in humans (see also ^8^). Thus, there is some evidence to suggest, in general terms, better performance for oppPE methods over B_0_ field maps, with nonlinear registration yielding poorer results (but better than no correction, and useful in cases where the additional scans required to perform the other methods above are not available).

This study considered only static geometric distortions in the GE EPI data caused by B_0_ inhomogeneity. If a subject moves during a scanning session, then the B_0_ inhomogeneities will not be stable over time, and geometric distortions will vary with head motion^7^, resulting in poorer correction based on static methods^10^. We conducted a second analysis (Analysis #2: pre-aligned data) in which all field map scans were first aligned to the magnitude portion of the B_0_ field map prior to distortion compensation. The results from this second analysis (Supplemental Figure 10) recapitulated the whole-brain findings from the main portion of our study (Figure 4), suggesting that differences in subject head motion over time may not explain the pattern of results we observed. Methods for dynamic distortion compensation (e.g., with different distortion fields calculated for each time point in an EPI time series) have also been proposed^14, 35, 37^, and may offer advantages in correcting time-varying geometric distortion, as compared to the static approaches considered here. However, to our knowledge, such dynamic distortion compensation methods are not currently implemented in the software packages that are most often used to pre-process brain imaging data.

## Acknowledgments

We thank Jesslyn (Li Shen) Chong, Tori Espensen-Sturges, Andrea N. Grant, Rohit S. Kamath, Timothy J. Lano, and Marisa J. Sanchez for their assistance with data collection. We also thank Bryon A. Mueller and Hannah R. Moser for help with data processing, and Essa Yacoub for supporting the design of the study.

This work was supported by funding from the National Institutes of Health (U01 MH108150). Salary support for MPS was provided by K01 MH120278. Salary support for CAO was provided by R01 MH111447. Support for MR scanning at the University of Minnesota Center for Magnetic Resonance Research was provided by P41 EB015894 and P30 NS076408. This work used tools from the University of Minnesota Clinical and Translational Science Institute that were supported by UL1 TR002494. The authors declare that they have no conflicts of interest with regard to the publication of this manuscript.

## Supplemental information

**Supplemental Figure 1.**
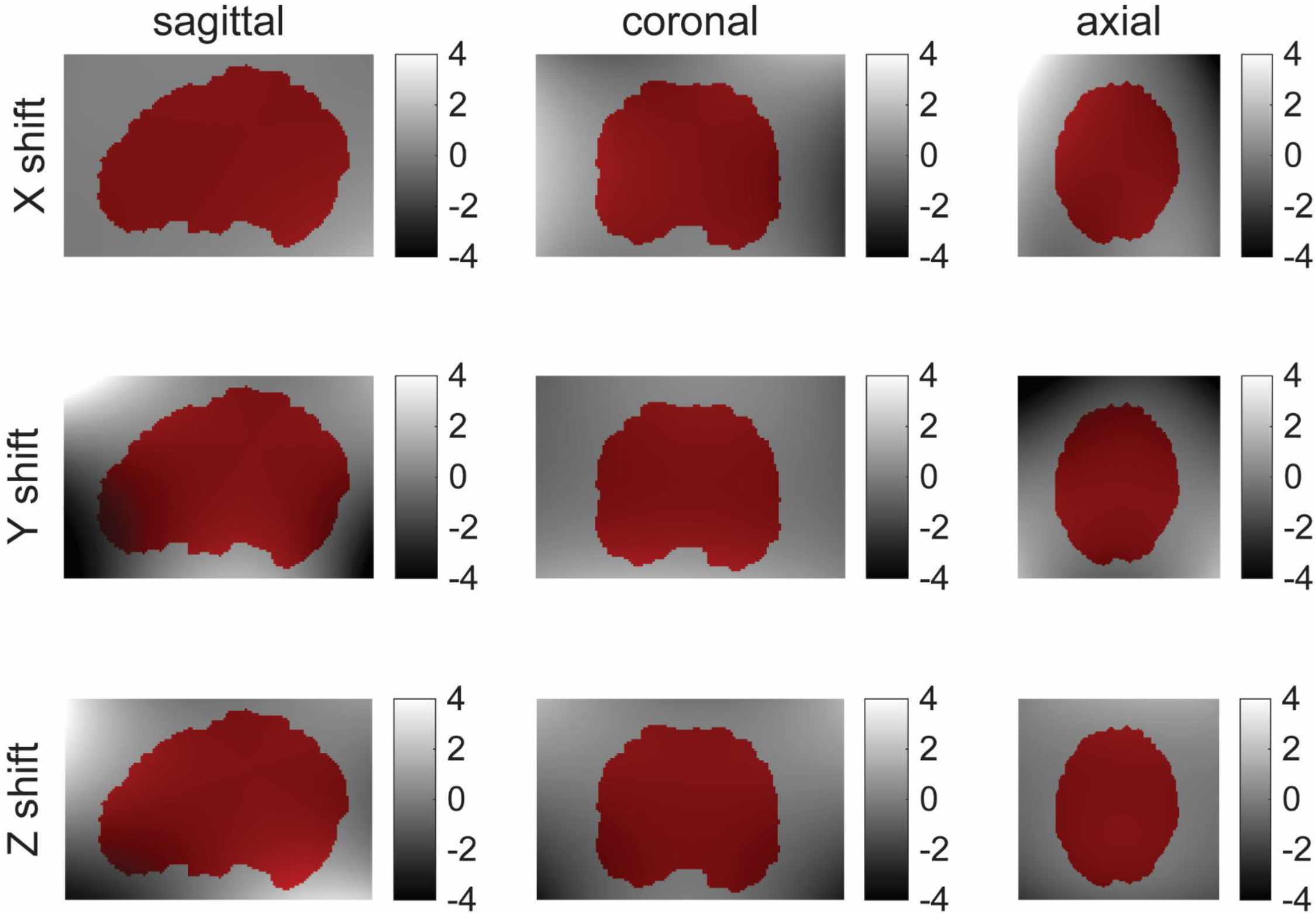
Example voxel shift images from gradient nonlinearity unwarping. Grayscale values show voxel shift in millimeters for each of 3 gradient directions (x, y, z) on separate rows. Columns show different views (left: sagittal, middle: coronal, right: axial). Data are from a single subject in Analysis #1 (main study). A partially transparent brain mask from the T_1_ anatomical scan is shown in red for reference. Notable distortions due to gradient nonlinearities are visible, especially near the edges of the brain and the field of view.

**Supplemental Table 1.**
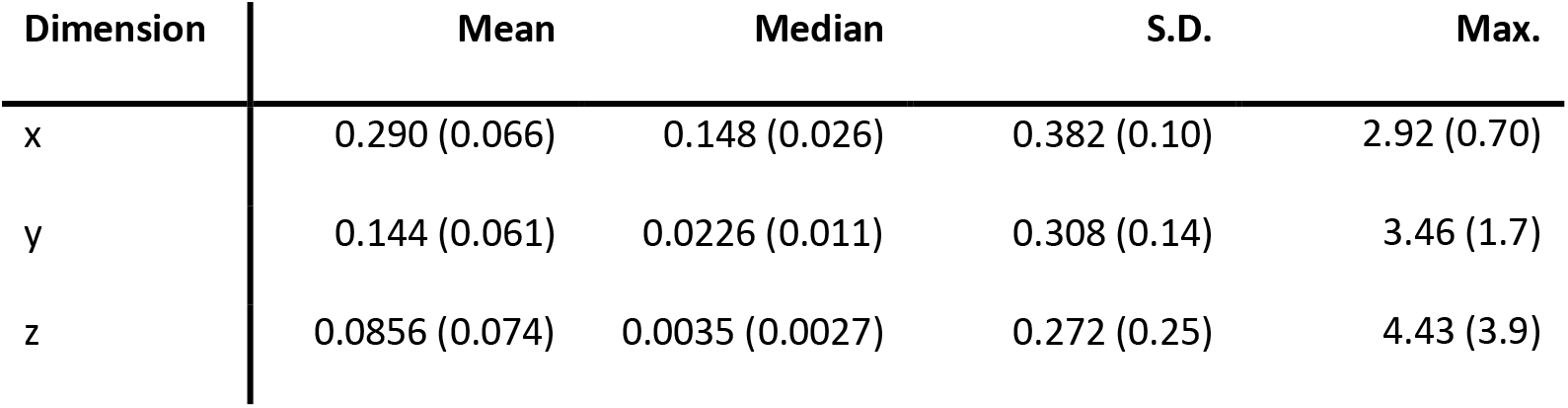
Summary of voxel shift data from gradient nonlinearity correction. Left column indicates the gradient direction (x, y, or z). For each of the 31 subjects, voxel shift values were obtained from *gradunwarp* for all voxels; the data shown here are restricted to voxels within the brain using a T_1_ anatomical mask. Note that the absolute value was taken for each voxel to permit averaging of shifts in the positive and negative directions. The mean, median, standard deviation (S.D.) and maximum (max.) voxel shift (in millimeters) were then calculated across voxels within each subject. The data shown here are the mean (S.D.) of each of these four metrics across all subjects.

**Supplemental Figure 2.**
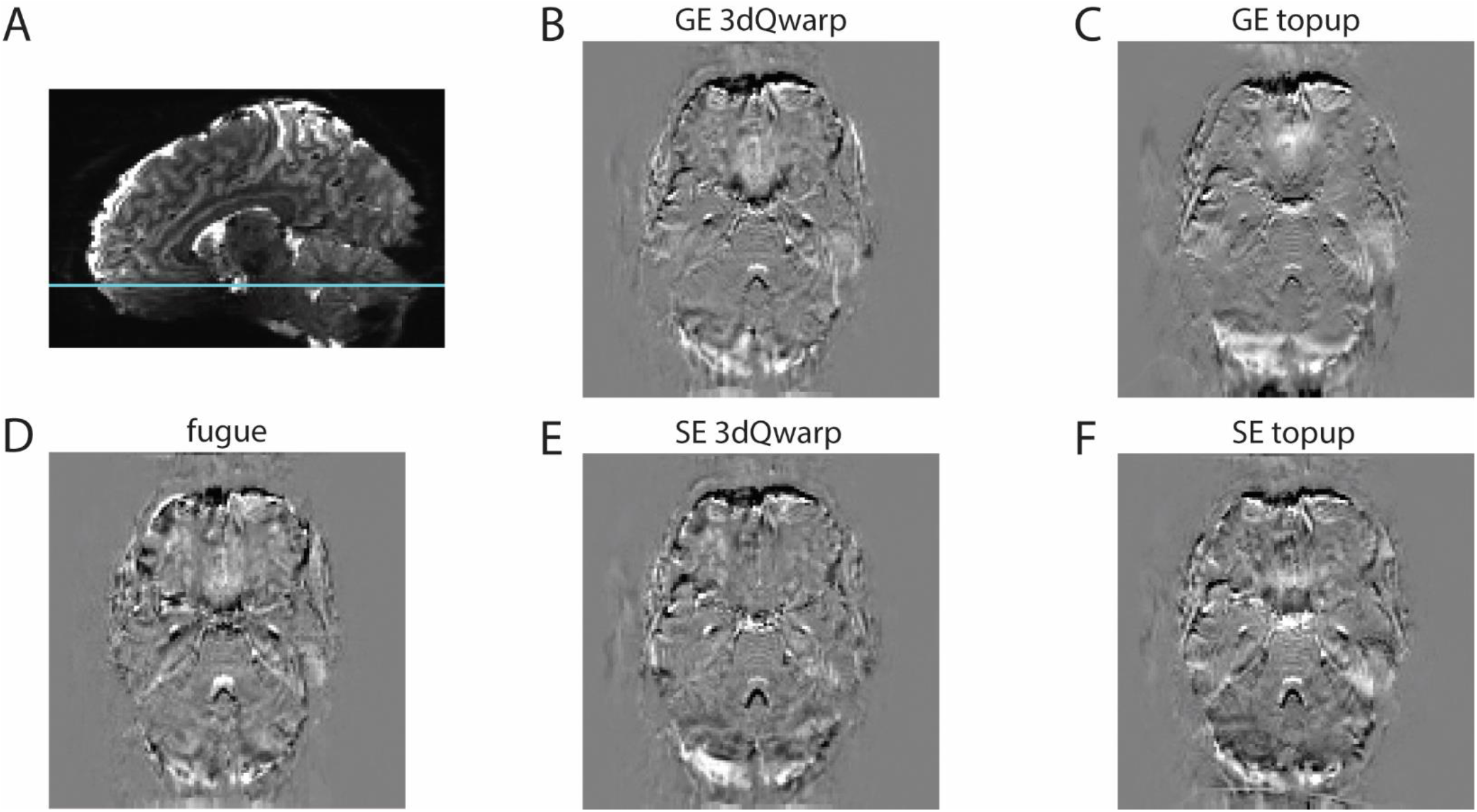
Difference between uncorrected and distortion-corrected data. **A**) Parasagittal section showing uncorrected EPI data from an example subject. Cyan line marks the position of axial sections in **B**-**D**. Distortion corrected GE EPI data were subtracted from the uncorrected data (i.e., uncorrected – corrected), following correction for geometric distortion using the following five correction methods: **B**) GE oppPE correction applied using AFNI’s *3dQwarp*, or **C**) FSL’s *topup*, **D**) B_0_ field map correction applied using FSL’s *fugue*, and **E**) SE oppPE correction applied using AFNI’s *3dQwarp*, or **F**) FSL’s *topup*. Brighter regions indicate greater signal in the uncorrected data, whereas darker regions indicate greater signal for the distortion corrected data. All brain images in **B**-**D** are examples from the same axial section in the same subject in Analysis #1 (main study), after gradient nonlinearity correction, and are scaled equally for comparison.

**Supplemental Figure 3.**
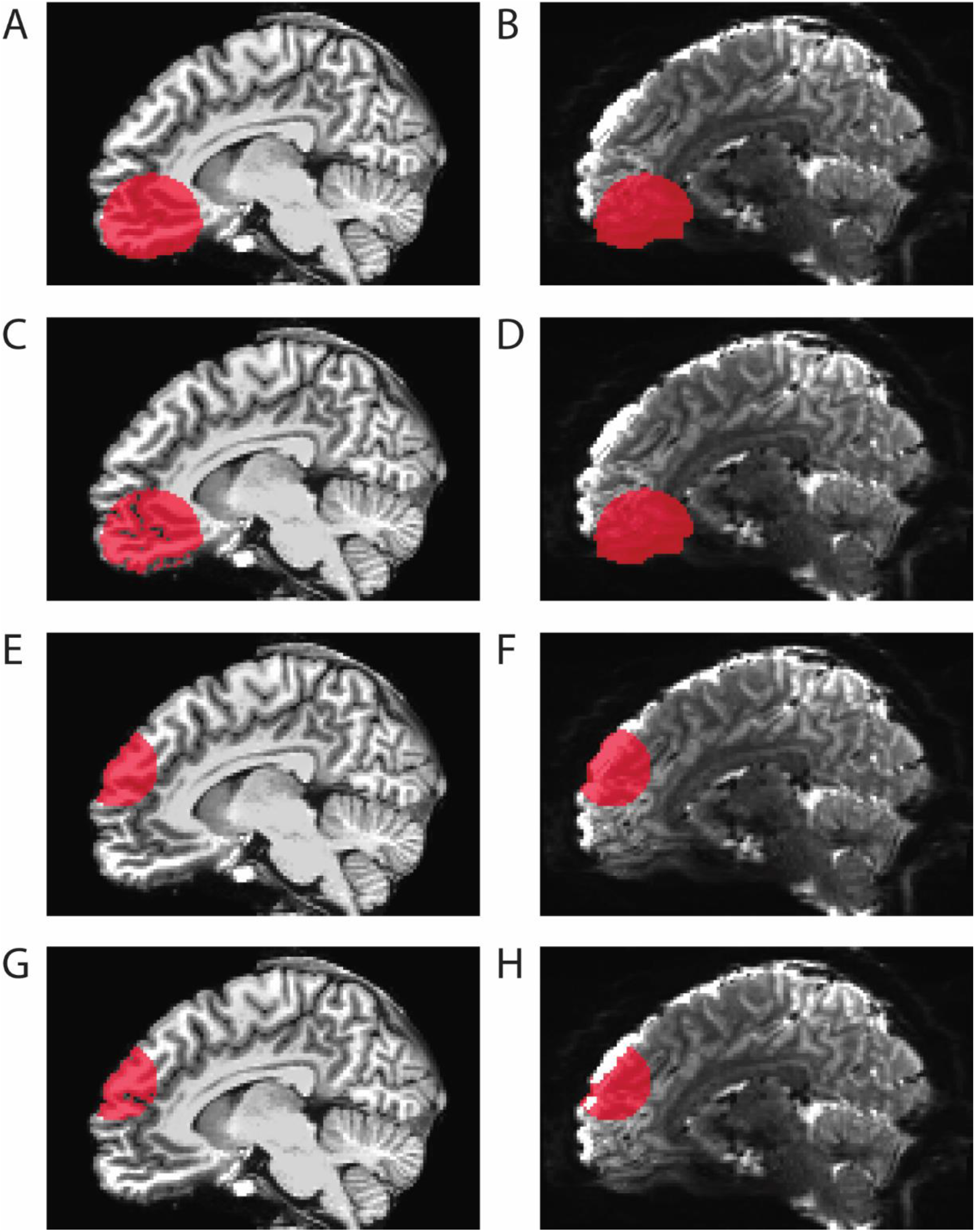
Regions of interest (ROIs). ROI analyses were performed in ventromedial prefrontal cortex (vmPFC; **A**-**D**) and dorsomedial prefrontal cortex (dmPFC; **C**-**H**). Red regions show example binary ROI masks for T_1_ anatomical (**A**, **C**, **E**, & **G**) and EPI (**B**, **D**, **F**, & **H**) scan data. **A** & **B** show whole-ROI masks for vmPFC (results shown in Figure 6A). **C** & **D** show CSF-excluded masks for vmPFC (results shown in Figure 6B). **E** & **F** show whole-ROI masks for dmPFC (results shown in Figure 7A). **G** & **H** show CSF-excluded masks for dmPFC (results shown in Figure 7B). All images show the same parasagittal section in the same example subject from the GE 3dQwarp condition in Analysis #1 (main study).

**Supplemental Figure 4.**
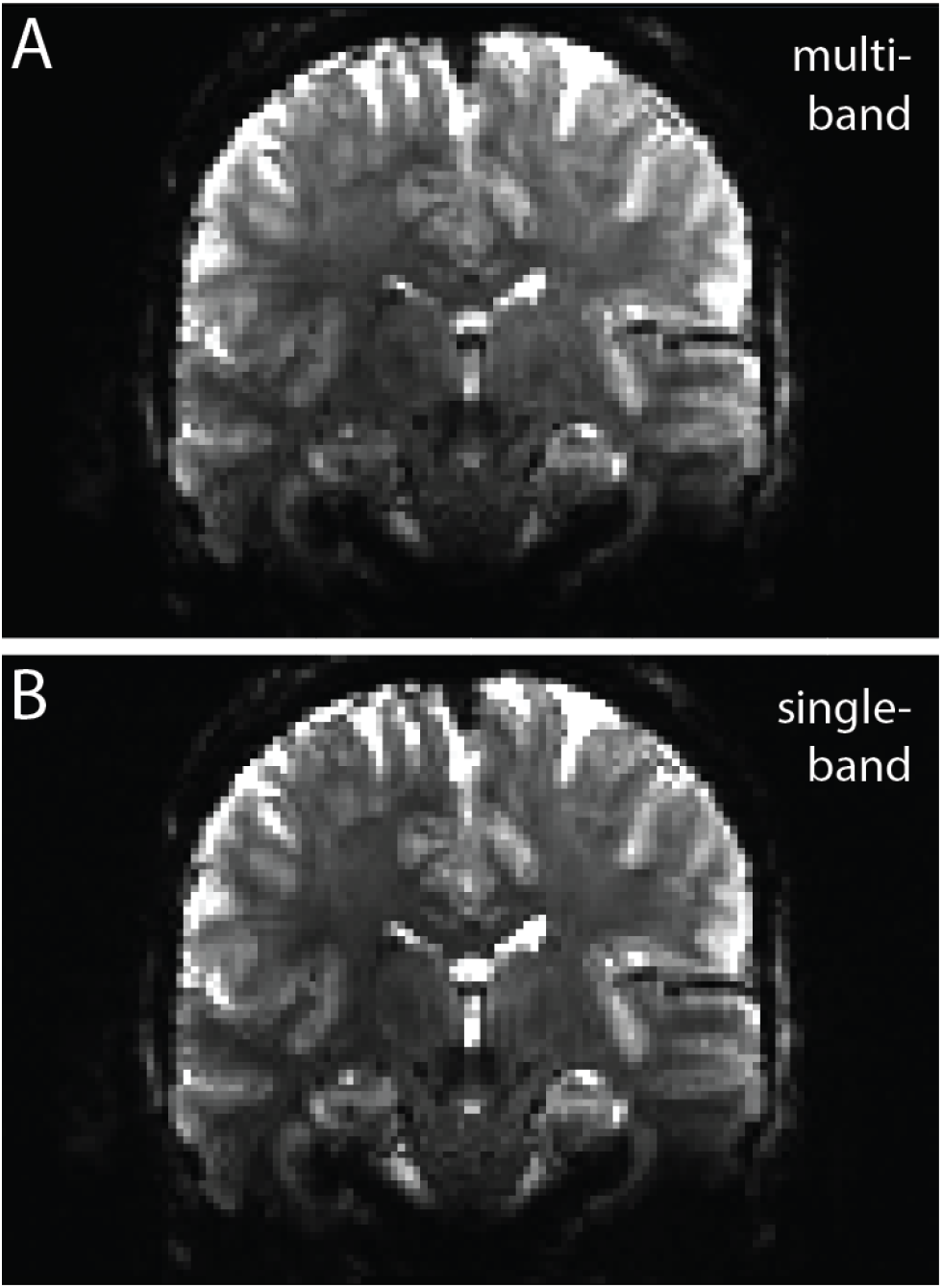
Single-band reference data. Panel **A** shows an example re-sliced coronal section from our standard multi-band GE EPI sequence, whereas **B** shows the single-band reference data from the same section in the same subject. Note the higher gray matter-white matter contrast for the single-band reference data in **B**.

**Supplemental Figure 5.**
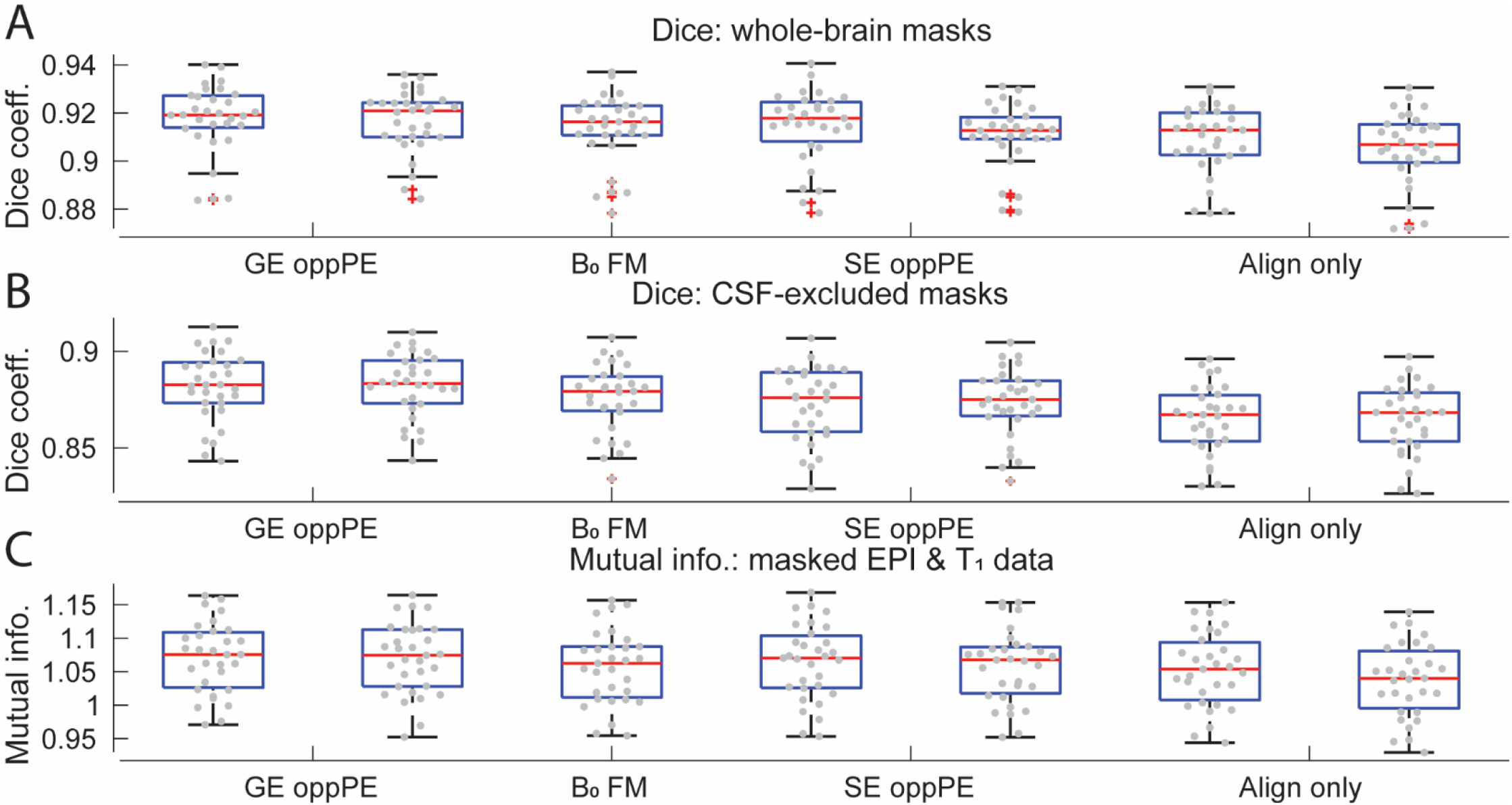
Box plots of the results from the main analysis. The order of conditions (left to right) is GE 3dQwarp, GE topup, fugue, SE 3dQwarp, SE topup, 6 param., 12 param. (same as in Figure 4). X-axis labels: GE oppPE = gradient echo opposite phase encoding field map, B_0_ FM = B_0_ field map, SE oppPE = spin echo opposite phase encoding field map, Align only = alignment-only (no explicit geometric distortion compensation). Red lines = median, blue boxes = interquartile range, whiskers = 1.5 × interquartile range, red pluses = points outside whiskers, gray dots = all data points.

**Supplemental Figure 6.**
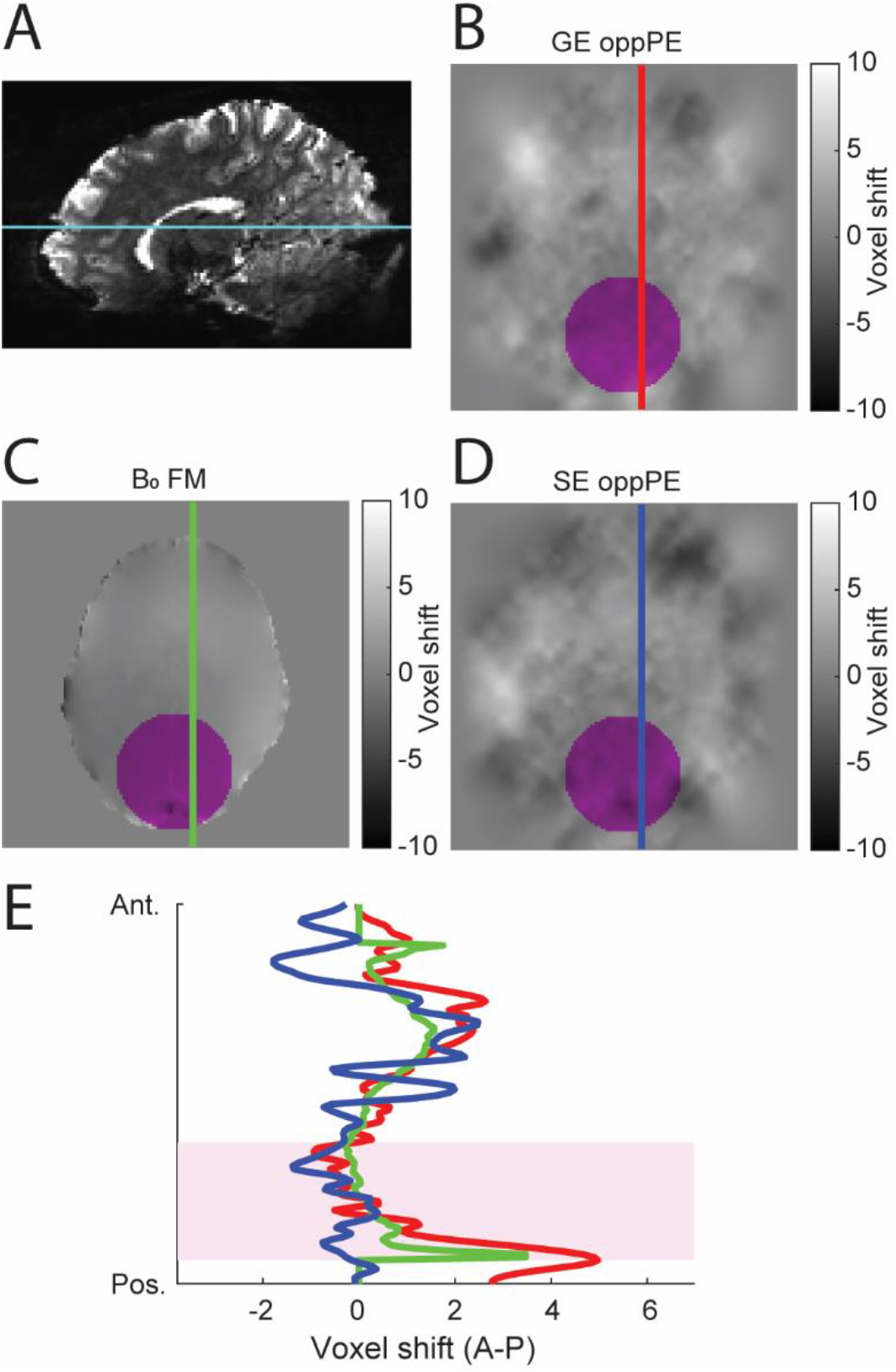
Posterior ROI voxel shift maps. Example voxel shift maps from the same ventral axial section (cyan line; **A**) in the same subject are shown for distortion compensation based on GE oppPE (**B**), B_0_ field map (**C**), and SE oppPE (**D**) correction methods. Color bar indicates voxel shift in the anterior-posterior (positive-negative) direction. Colored vertical lines indicate the positions of the voxel shift data plotted in **E**. Magenta regions in **B**-**E** show the posterior ROI position. GE and SE oppPE corrections in these examples were performed with AFNI’s *3dQwarp*. Data are from Analysis #1 (main study). Differences in voxel shift maps between methods are visible in posterior cortical regions.

**Supplemental Figure 7.**
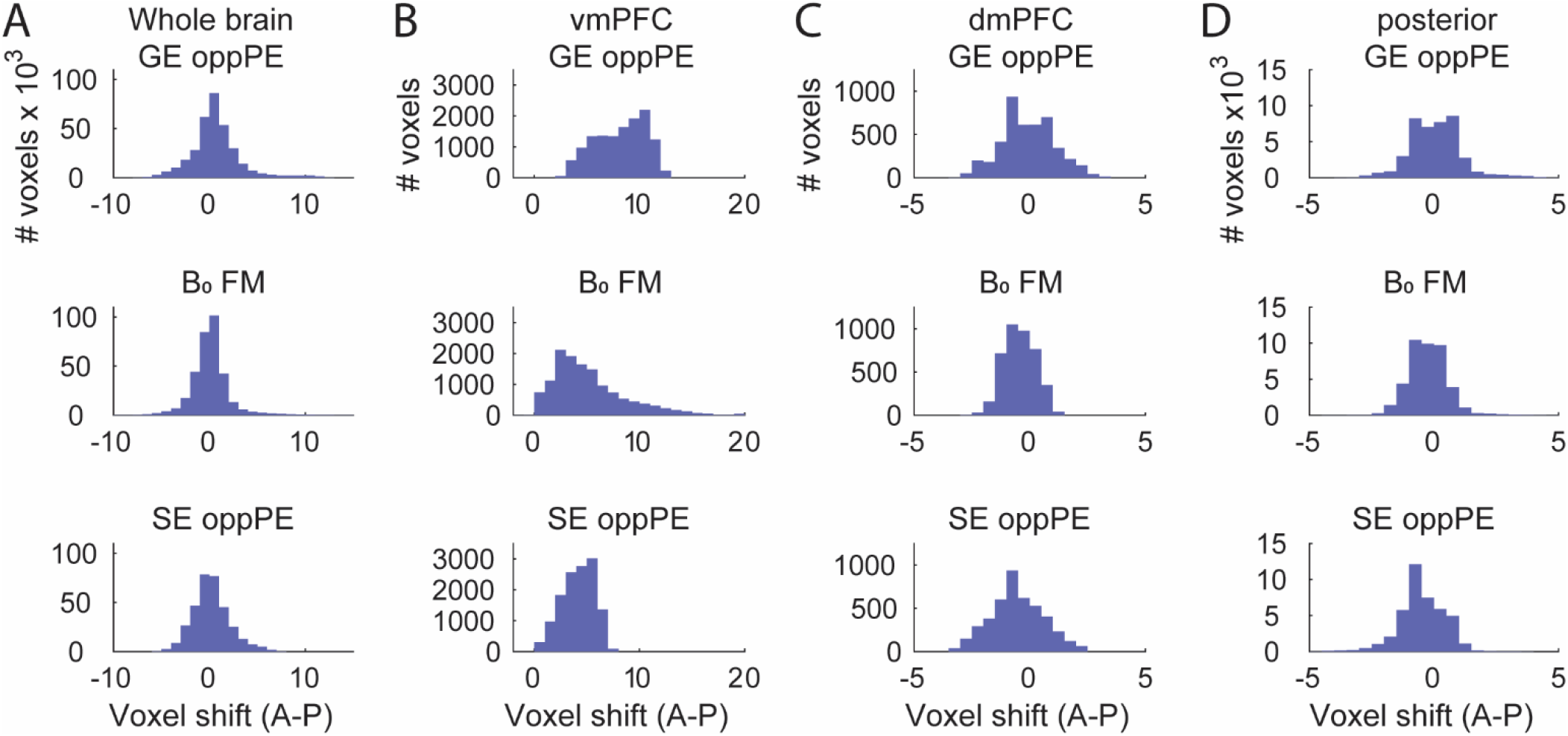
Example distortion compensation voxel shift histograms. Voxel shifts in anterior-posterior (positive-negative) direction are shown for the whole brain (**A**), as well as vmPFC (**B**), dmPFC (**C**), and posterior (**D**) ROIs. Data obtained using different methods for correcting geometric distortion due to B_0_ inhomogeneity are shown in each row. Top row shows GE oppPE data, middle row shows B_0_ field map, bottom row shows SE oppPE. Note that the x- and y-axes differ across columns. Data are from a single subject in Analysis #1 (main study), and are restricted to voxels within the T_1_ anatomical brain mask. Subtle differences between methods (rows) are visible across all regions, with more dramatic differences in vmPFC (**B**).

**Supplemental Figure 8.**
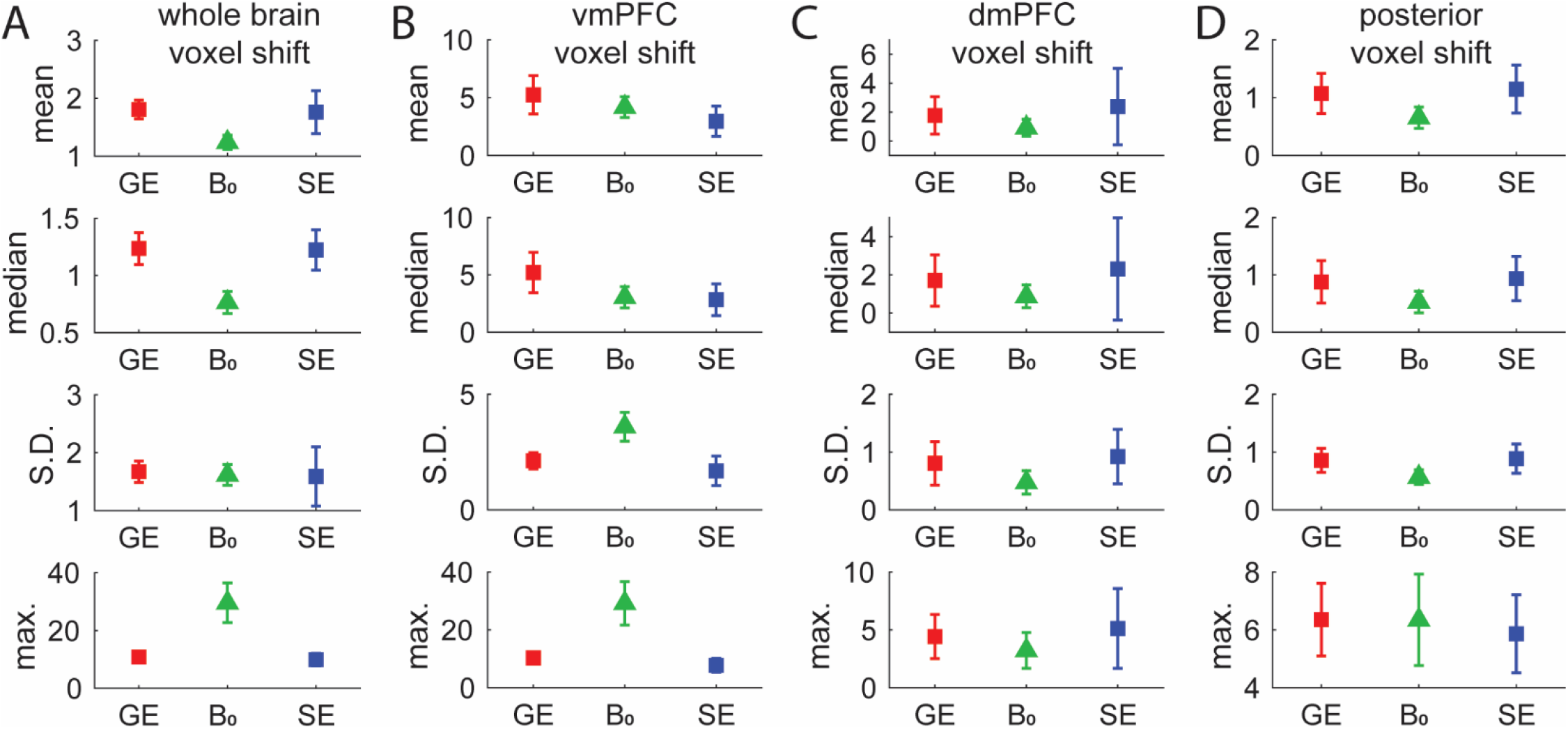
Voxel shift summary data from B_0_ inhomogeneity distortion compensation. Each column shows summary data for different brain regions (**A**: whole brain, **B**: vmPFC, **C**: dmPFC, **D**: posterior). For each of the 31 subjects, voxel shift values were obtained for all voxels within the brain using the T_1_ anatomical mask. ROI analyses (**B-D**) were further restricted using the relevant ROI mask. Note that the absolute value was taken for each voxel to allow averaging of shifts in the positive and negative directions. The mean, median, standard deviation (S.D.) and maximum (max.) voxel shift (in millimeters) were then calculated across voxels within each subject. Symbols show mean, error bars show S.D. of these four metrics across all subjects. GE = GE oppPE data from *3dQwarp*, B_0_ = B_0_ field map data from *fugue*, SE = SE oppPE data from *3dQwarp*. These data are from Analysis #1 (main study), and show notable variability in the voxel shifts for correcting geometric distortion due to B_0_ inhomogeneity across different methods and brain regions.

**Supplemental Figure 9.**
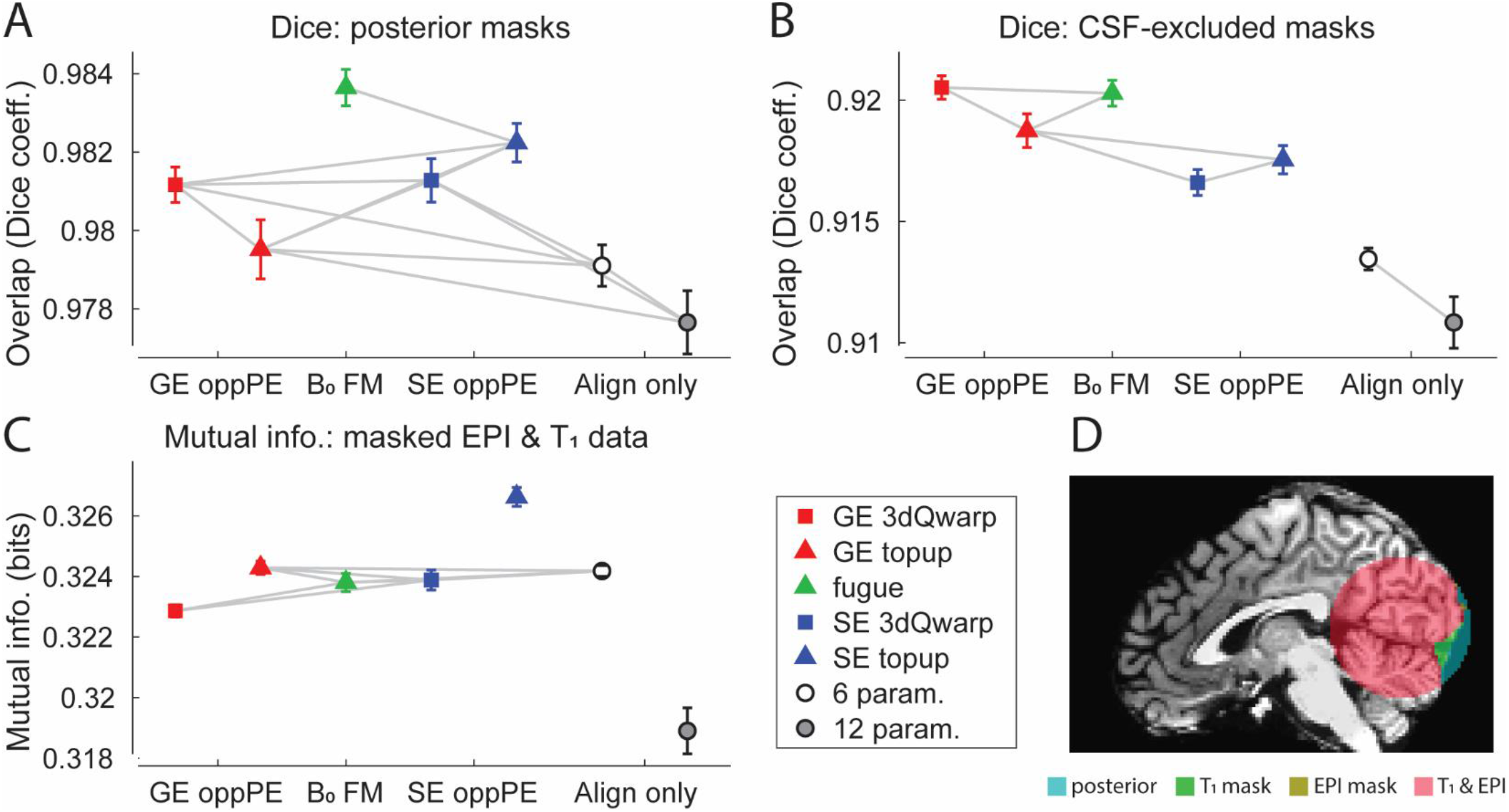
Results in posterior brain regions. **A**) Overlap (Dice coefficient) between GE EPI and T_1_ binary masks restricted to a posterior ROI across different distortion compensation methods. Gray lines indicate conditions that do <u>not</u> differ significantly (post-hoc paired *t*-tests, threshold *p* < 0.05, FDR corrected). X-axis labels: GE oppPE = gradient echo opposite phase encoding field map (red), B_0_ FM = B_0_ field map (green), SE oppPE = spin echo opposite phase encoding field map (blue), Align only = alignment-only (no explicit geometric distortion compensation). **B**) Same, but for binary masks in posterior brain areas with regions of cerebrospinal fluid (CSF) excluded, following tissue segmentation. **C**) Mutual information between GE EPI and T_1_ scan data in the posterior ROI. Squares show data corrected using AFNI, triangles show data from FSL, circles show alignment-only data. Error bars are *SEM* calculated within subjects^1^. **D**) Example T_1_ anatomical image with overlaid posterior masks: whole-ROI (teal), T_1_ (green), EPI (yellow), and T_1_-EPI overlap (red). The Dice coefficient analysis in **A** quantified the agreement between EPI and T_1_ masks (i.e., red versus green and yellow), whereas **B** did the same after excluding CSF regions. T_1_-EPI agreement in posterior regions was similar across distortion compensation methods, and tended to be higher than in the alignment-only conditions.

**Supplemental Figure 10.**
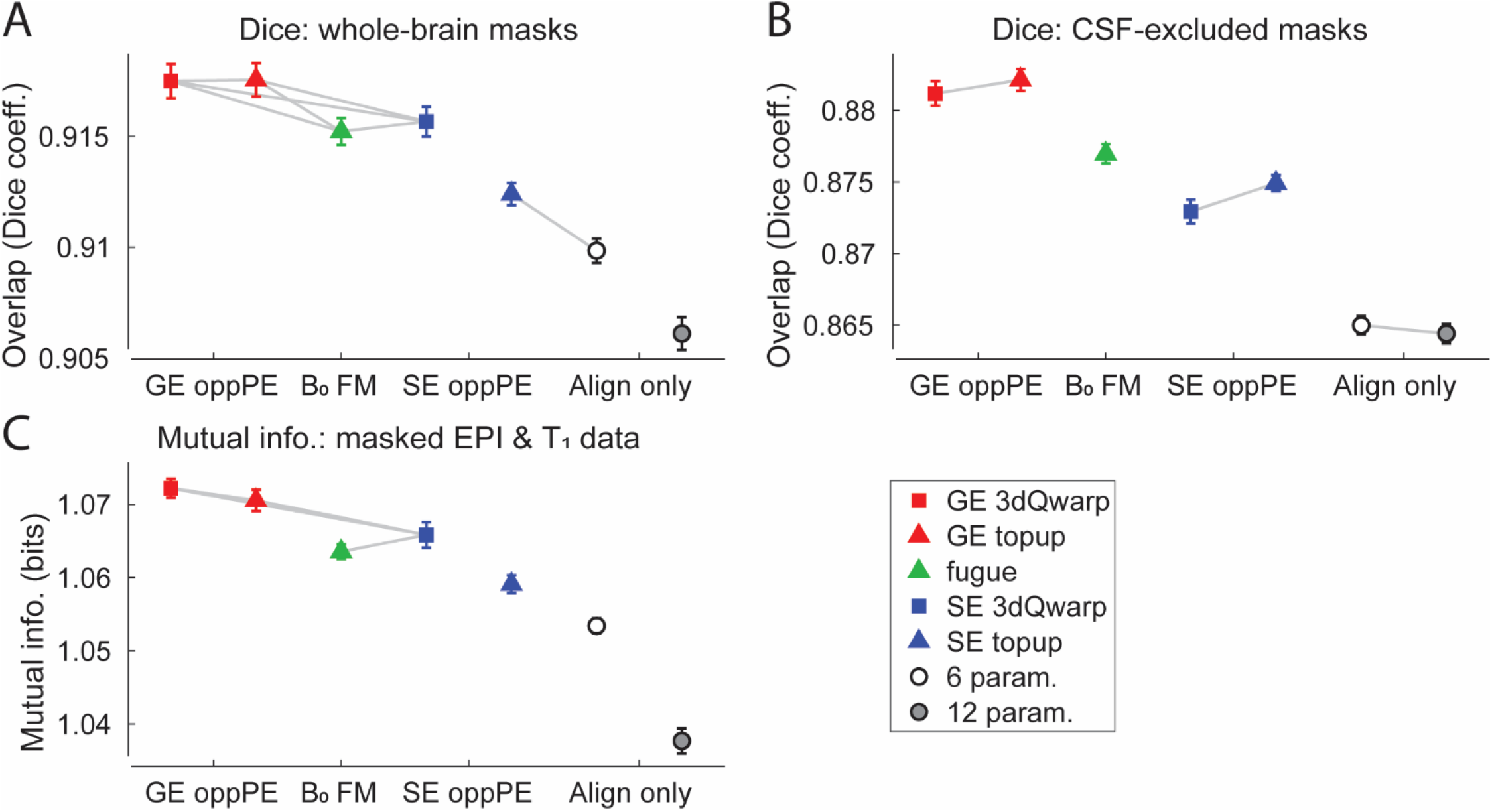
Results for data aligned to B_0_ field map from Analysis #2. **A**) Overlap (Dice coefficient) between GE EPI and T_1_ brain masks across different distortion compensation methods. Gray lines indicate conditions that do <u>not</u> differ significantly (post-hoc paired *t*-tests, threshold *p* < 0.05, FDR corrected). X-axis labels: GE oppPE = gradient echo opposite phase encoding field map (red), B_0_ FM = B_0_ field map (green), SE oppPE = spin echo opposite phase encoding field map (blue), Align only = alignment-only (no explicit geometric distortion compensation). **B**) Same, but for binary masks with regions of cerebrospinal fluid (CSF) excluded, following tissue segmentation. **C**) Mutual information between GE EPI and T_1_ scan data within whole-brain masks. Squares show data corrected using AFNI, triangles show data from FSL, circles show alignment-only data. Error bars are *SEM* calculated within subjects^1^. Aligning all scans to the magnitude portion of the B_0_ field map did not substantially alter our pattern of results (compare with data from the main analysis in Figure 4), suggesting that differences in head motion across the scanning session did not dramatically affect our findings.

**Supplemental Figure 11.**
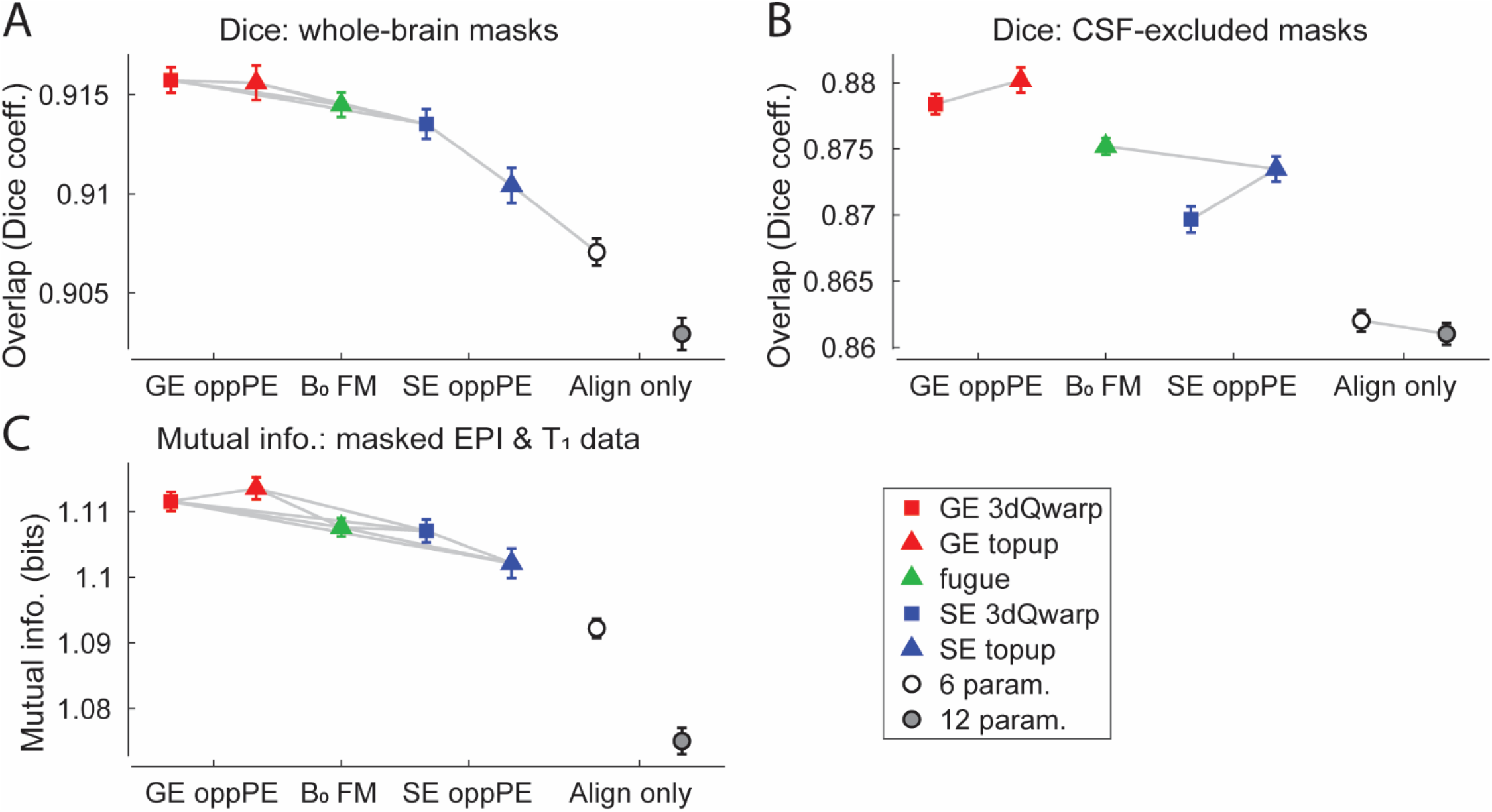
Results for single-band reference data from Analysis #3. **A**) Overlap (Dice coefficient) between GE EPI and T_1_ brain masks across different distortion compensation methods. Gray lines indicate conditions that do <u>not</u> differ significantly (post-hoc paired *t*-tests, threshold *p* < 0.05, FDR corrected). X-axis labels: GE oppPE = gradient echo opposite phase encoding field map (red), B_0_ FM = B_0_ field map (green), SE oppPE = spin echo opposite phase encoding field map (blue), Align only = alignment-only (no explicit geometric distortion compensation). **B**) Same, but for binary masks with regions of cerebrospinal fluid (CSF) excluded, following tissue segmentation. **C**) Mutual information between GE EPI and T_1_ scan data within whole-brain masks. Squares show data corrected using AFNI, triangles show data from FSL, circles show alignment-only data. Error bars are *SEM* calculated within subjects^1^. Using the single-band reference data for alignment and segmentation did not dramatically alter the pattern of results (compare with data from the main analysis in Figure 4), suggesting that our findings did not depend strongly on image contrast.

**Supplemental Figure 12.**
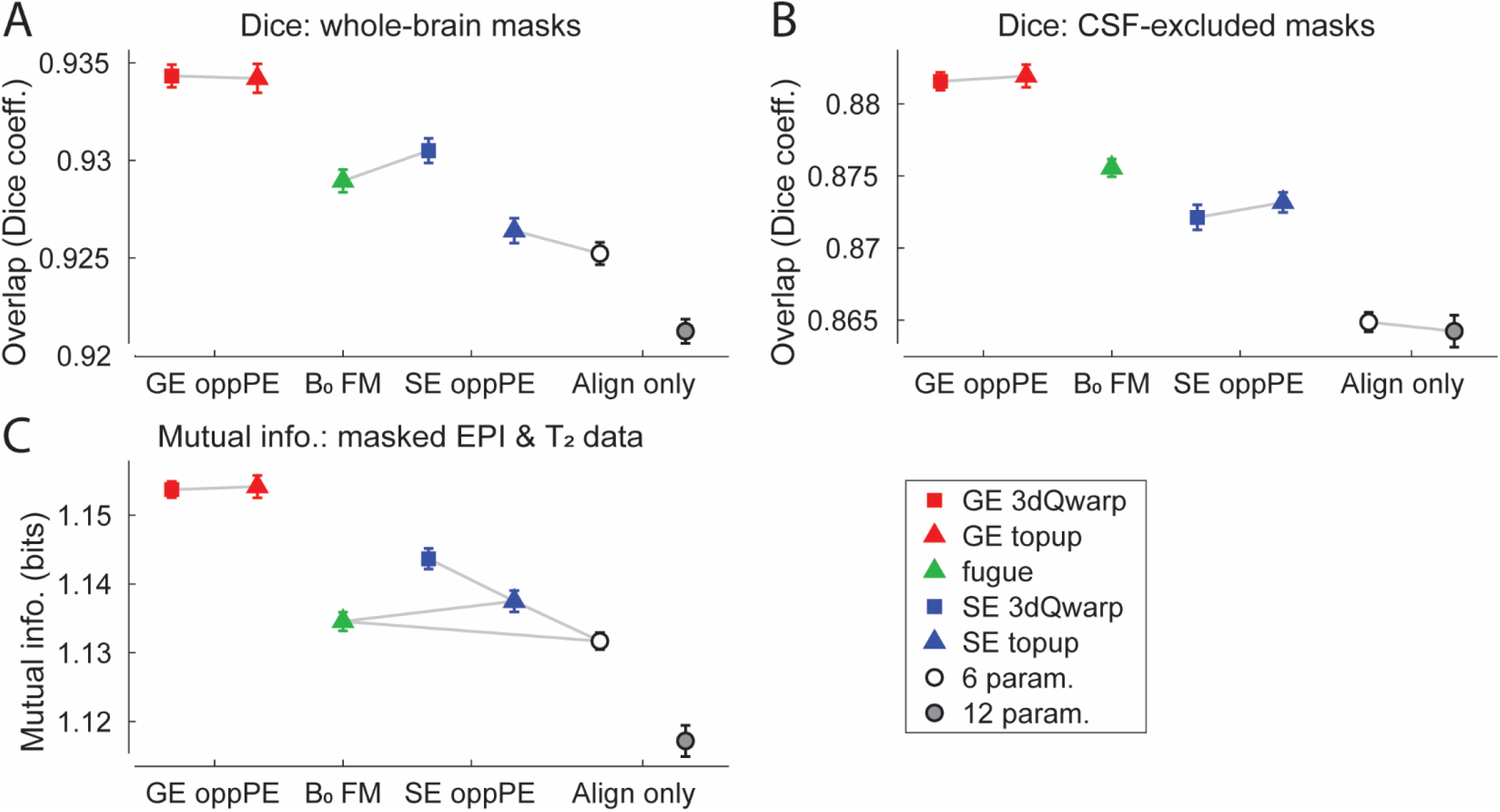
Results using the T_2_ anatomical scan as a reference, from Analysis #4. **A**) Overlap (Dice coefficient) between GE EPI and T_2_ brain masks across different distortion compensation methods. Gray lines indicate conditions that do <u>not</u> differ significantly (post-hoc paired *t*-tests, threshold *p* < 0.05, FDR corrected). X-axis labels: GE oppPE = gradient echo opposite phase encoding field map (red), B_0_ FM = B_0_ field map (green), SE oppPE = spin echo opposite phase encoding field map (blue), Align only = alignment-only (no explicit geometric distortion compensation). **B**) Same, but for binary masks with regions of cerebrospinal fluid (CSF) excluded, following tissue segmentation. **C**) Mutual information between GE EPI and T_2_ scan within the respective whole-brain masks. Squares show data corrected using AFNI, triangles show data from FSL, circles show alignment-only data. Error bars are *SEM* calculated within subjects^1^. These results are highly similar to those we found using a T_1_ anatomical reference scan in our main analysis (Figure 4), suggesting that the choice of reference scan has little effect on our findings.

**Supplemental Figure 13.**
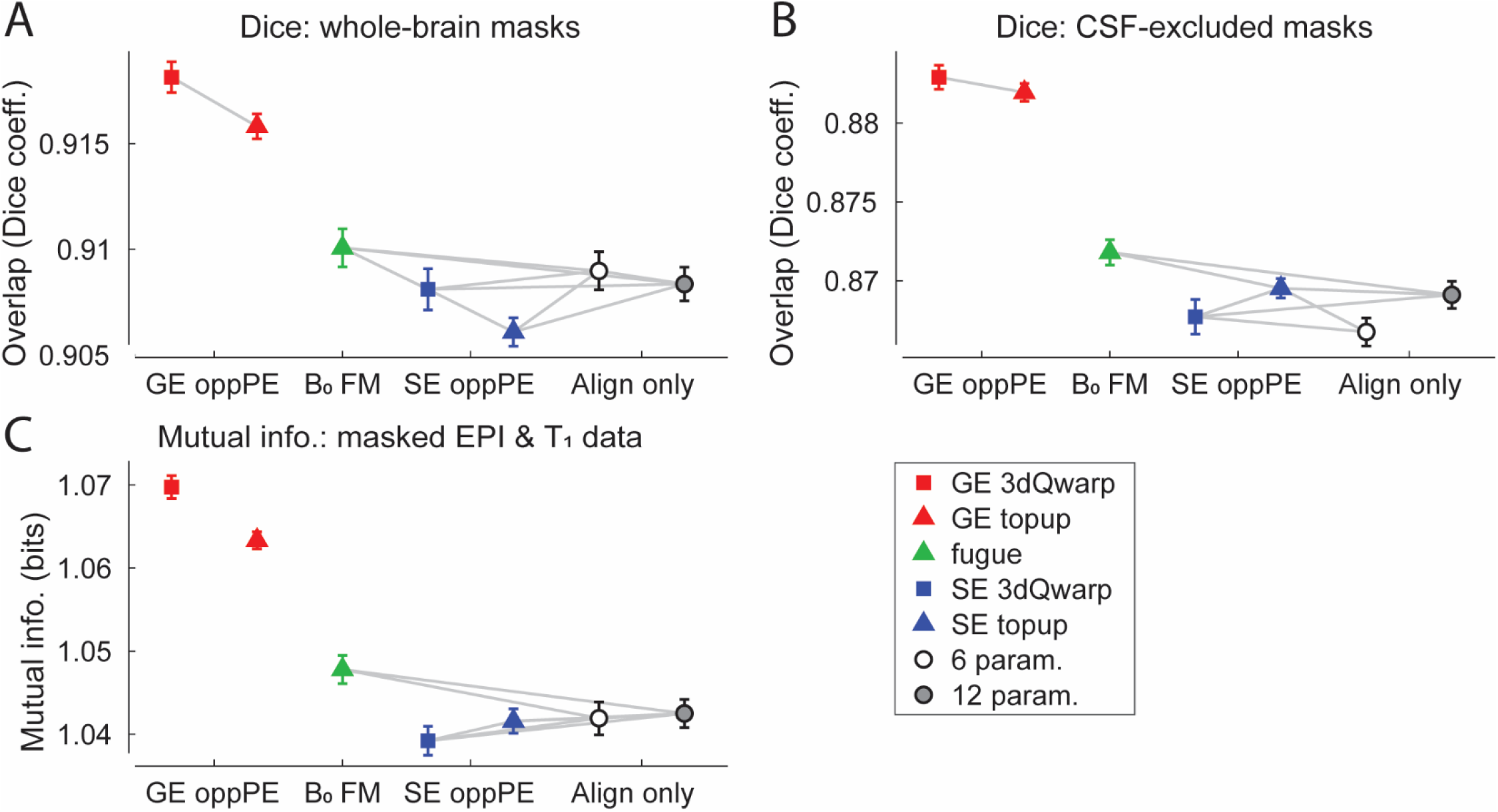
Results using posterior-anterior GE data from Analysis #5. **A**) Overlap (Dice coefficient) between GE EPI and T_1_ brain masks across different distortion compensation methods. Gray lines indicate conditions that do <u>not</u> differ significantly (post-hoc paired *t*-tests, threshold *p* < 0.05, FDR corrected). X-axis labels: GE oppPE = gradient echo opposite phase encoding field map (red), B_0_ FM = B_0_ field map (green), SE oppPE = spin echo opposite phase encoding field map (blue), Align only = alignment-only (no explicit geometric distortion compensation). **B**) Same, but for binary masks with regions of cerebrospinal fluid (CSF) excluded, following tissue segmentation. **C**) Mutual information between GE EPI and T_1_ scan data within whole-brain masks. Squares show data corrected using AFNI, triangles show data from FSL, circles show alignment-only data. Error bars are *SEM* calculated within subjects^1^. These results are generally similar to the pattern we observed using GE EPI data with an AP PE direction (from Analysis #2: pre-aligned data; Supplemental Figure 10), suggesting that the choice of AP versus PA phase encoding direction for our GE EPI data has relatively little impact on our results.

**Supplemental Figure 14.**
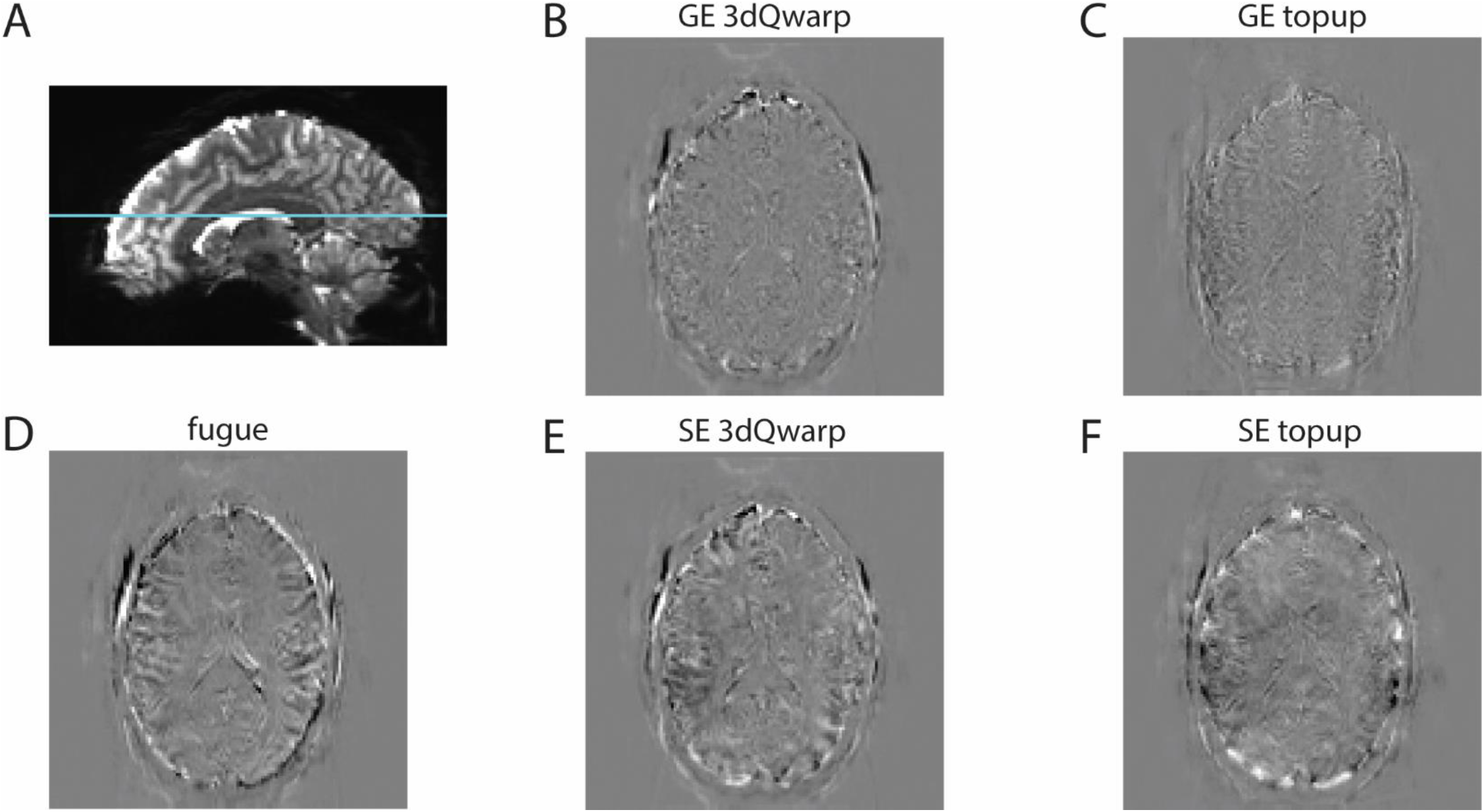
Example data showing the difference between AP and PA data after distortion compensation. **A**) Parasagittal section showing EPI data from an example subject. Cyan line marks the position of axial sections in **B-D**. GE EPI data with a PA PE direction were subtracted from corresponding AP data (i.e., AP – PA), following the correction of each for geometric distortion using the following five correction methods: **B**) GE oppPE correction applied using AFNI’s *3dQwarp*, or **C**) FSL’s *topup*, **D**) B_0_ field map correction applied using FSL’s *fugue*, and **E**) SE oppPE correction applied using AFNI’s *3dQwarp*, or **F**) FSL’s *topup*. Brighter regions indicate greater signal in the AP data, whereas darker regions indicate greater signal for the PA data. This subtraction method has previously been used to assess residual error following distortion compensation^33^. All brain images are examples from the same axial section in the same subject, with the same arbitrary scaling. AP data are from Analysis #2 (pre-aligned data), while PA data are from Analysis #5 (posterior-anterior PE data). Prior to distortion compensation, data in both analyses were aligned to the magnitude portion of the B_0_ field map, to mitigate possible effects of head motion between AP and PA scan acquisitions. The higher contrast (i.e., residual error) in the B_0_ field map and SE oppPE conditions (**D-F**) as compared to the GE oppPE data (**B** & **C**) suggests the AP and PA data were more similar for the GE oppPE analyses. Lower residual error is consistent with superior distortion compensation for the GE oppPE methods.

**Supplemental Figure 15.**
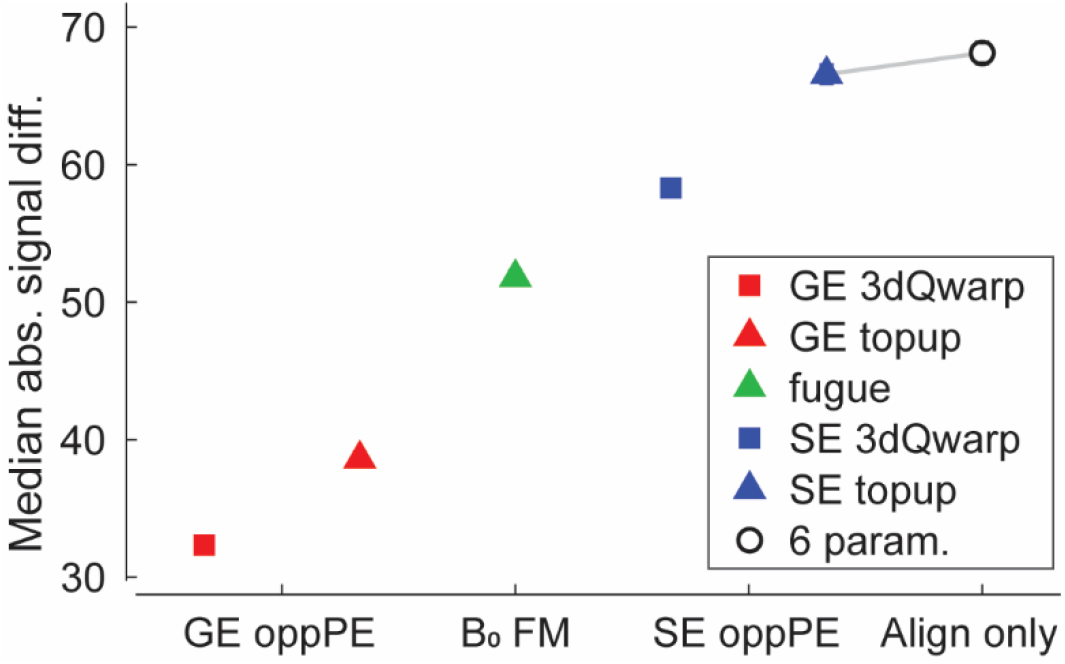
Residual error for AP minus PA data. Residual error was calculated by taking the difference between AP (Analysis #2: pre-aligned data) and PA (Analysis #5: posterior-anterior PE data) EPI data, as in Supplemental Figure 14. Data were masked with a summed whole-brain mask (AP + PA), the absolute value of each voxel intensity was taken (to allow averaging of positive and negative values), and then the median was calculated across voxels within each subject. Gray lines indicate conditions that do <u>not</u> differ significantly (post-hoc paired *t*-tests, threshold *p* < 0.05, FDR corrected). X-axis labels: GE oppPE = gradient echo opposite phase encoding field map (red), B_0_ FM = B_0_ field map (green), SE oppPE = spin echo opposite phase encoding field map (blue), Align only = alignment-only (no explicit geometric distortion compensation). Squares show data corrected using AFNI, triangles show data from FSL, circles show alignment-only data. Symbols are the mean across subjects. Error bars (generally smaller than symbols) are *SEM* calculated within subjects^1^. GE oppPE methods showed the lowest residual error (signal difference), consistent with superior distortion compensation of our GE EPI data.

**Supplemental Figure 16.**
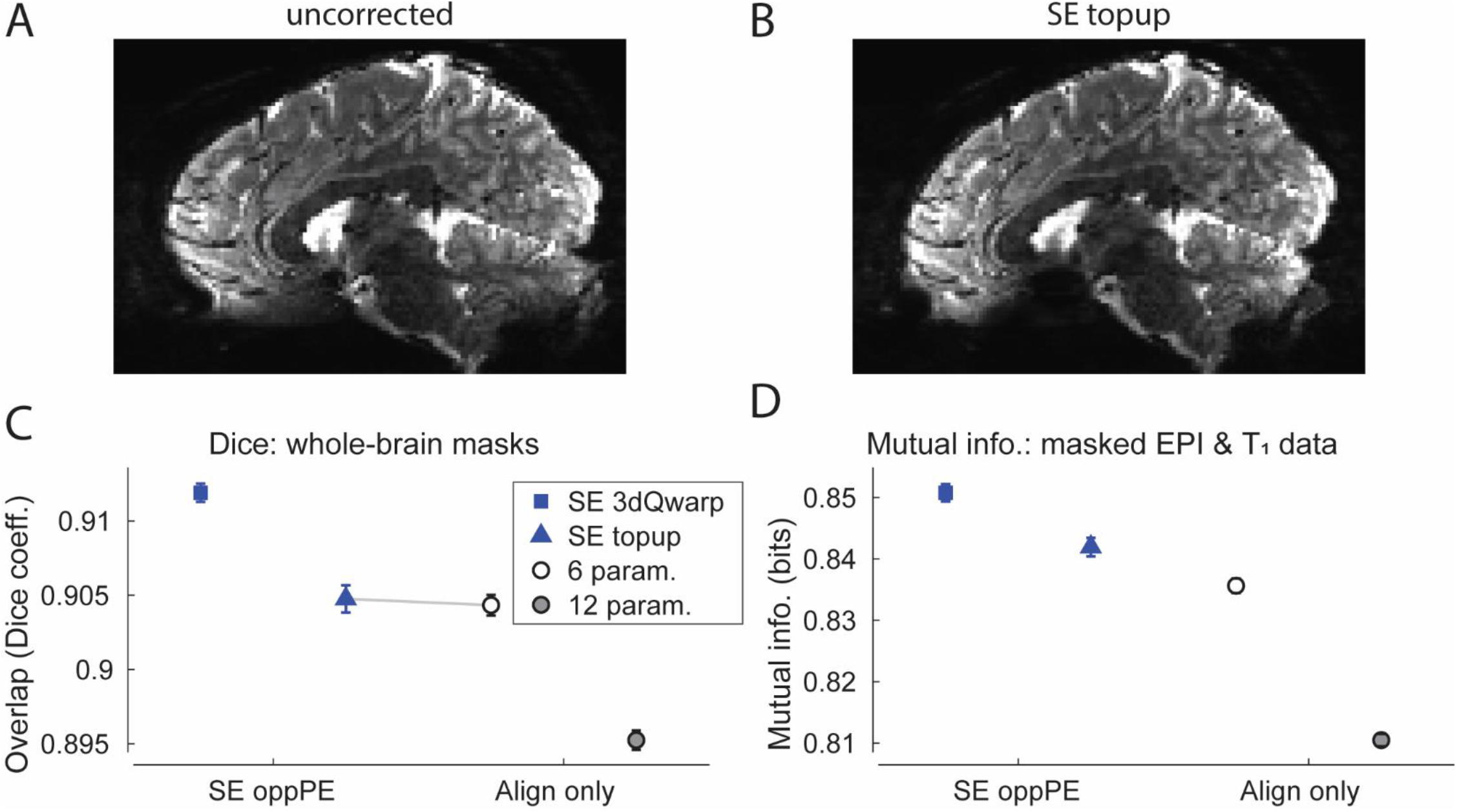
Young Adult HCP results from Analysis #6. Sagittal section from a single example subject, showing unprocessed (**A**) 7T GE EPI data, and the same section after SE oppPE distortion compensation within our pipeline via FSL’s *topup* (**B**).

**C**) Overlap (Dice coefficient) between GE EPI and T_1_ whole-brain mask data, across different analysis conditions. Gray line indicates conditions that do <u>not</u> differ significantly (post-hoc paired *t*-tests, threshold *p* < 0.05, FDR corrected). X-axis labels: SE oppPE = spin echo opposite phase encoding field map (blue), Align only = alignment-only (no explicit geometric distortion compensation). **D**) Mutual information between GE EPI and T_1_ scan data, within whole-brain masks. Squares show data corrected using AFNI, triangles show data from FSL, circles show alignment-only data. Error bars are *SEM* calculated within subjects^1^. These results show comparable agreement between T_1_ and EPI data to the findings from our own dataset (Figure 4). SE oppPE distortion correction (blue) increased correspondence between EPI and T_1_ anatomical data, compared to alignment-only methods.

**Supplemental Figure 17.**
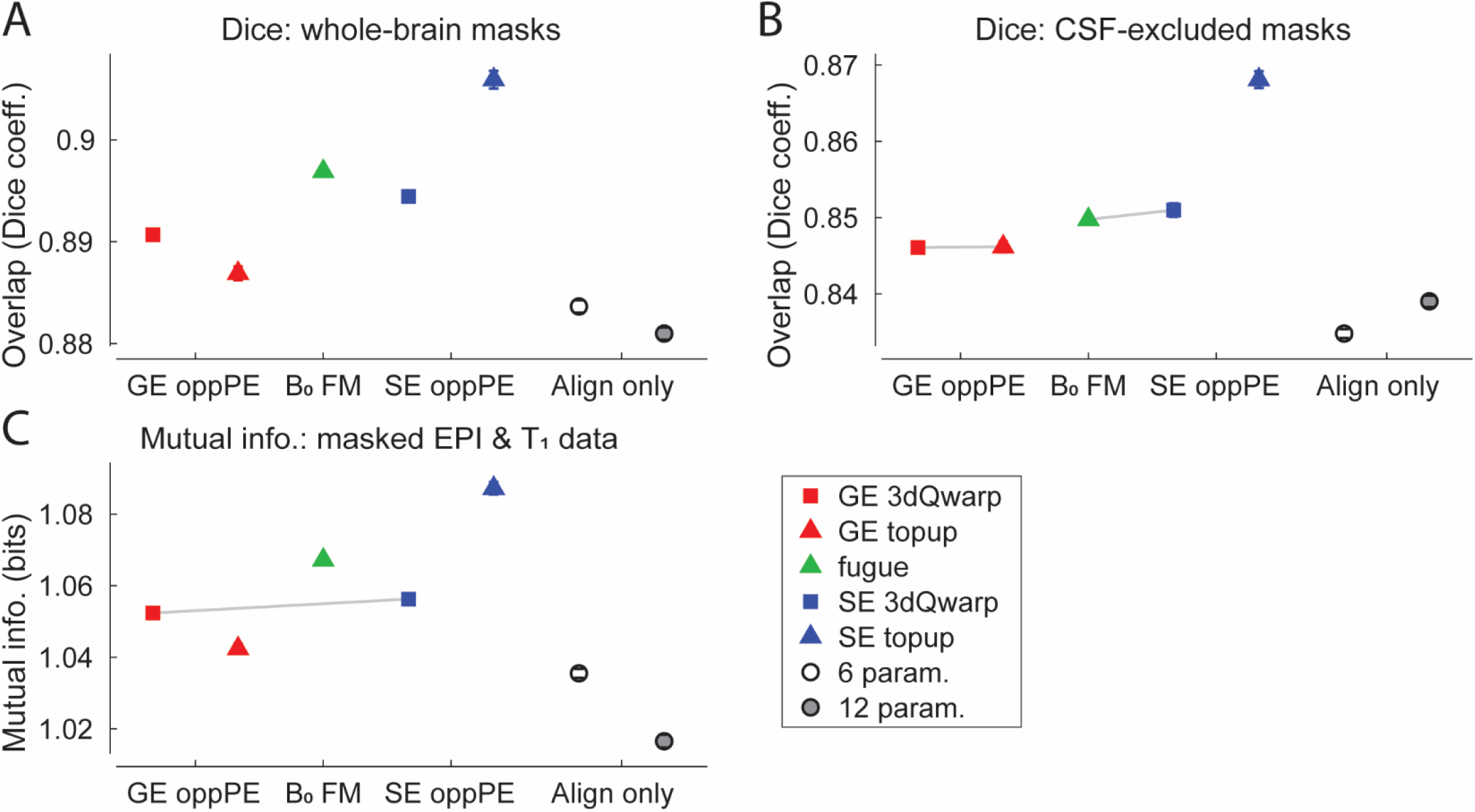
Results when correcting SE data from Analysis #7. **A**) Overlap (Dice coefficient) between SE EPI and T_1_ brain masks across different distortion compensation methods. Gray lines indicate conditions that do <u>not</u> differ significantly (post-hoc paired *t*-tests, threshold *p* < 0.05, FDR corrected). X-axis labels: GE oppPE = gradient echo opposite phase encoding field map (red), B_0_ FM = B_0_ field map (green), SE oppPE = spin echo opposite phase encoding field map (blue), Align only = alignment-only (no explicit geometric distortion compensation). **B**) Same, but for binary masks with regions of cerebrospinal fluid (CSF) excluded, following tissue segmentation. **C**) Mutual information between SE EPI and T_1_ scan data within the respective whole-brain masks. Squares show data corrected using AFNI, triangles show data from FSL, circles show alignment-only data. Error bars are *SEM* calculated within subjects^1^. Across the three metrics, SE oppPE methods (blue) tended to yield superior EPI-T_1_ alignment, as compared to GE oppPE field maps (red), with B_0_ field map correction via *fugue* showing intermediate performance. When considered along with our other results, this suggests that matching the type of field map scan to the EPI data to-be corrected (i.e., GE-to-GE or SE-to-SE) yields superior distortion compensation.

